# Bacterial RNA polymerase exemplifies a general physical mechanism for accelerating protein-DNA association

**DOI:** 10.64898/2026.07.21.739884

**Authors:** Francis Rivera, Alex Turek, Wencheng Ji, Matt Copeland, Larry Friedman, Fiona Stewart, Johnson Chung, Ariel Amir, Jane Kondev, Jeff Gelles

**Author notes:** To whom correspondence may be addressed: Jane Kondev, Dept. of Physics, MS 057, Brandeis University, P.O. Box 549110, Waltham, MA, USA, 02454., Jeff Gelles, Dept. of Biochemistry, MS 009, Brandeis University, P.O. Box 549110, Waltham, MA, USA, 02454.

## Abstract

Bacterial RNA polymerases (RNAPs) have two flexibly tethered α subunit C-terminal domains (α-CTDs) that bind DNA. Interaction between α-CTDs and some promoter DNA motifs is known to accelerate transcription initiation, but the physical mechanism by which it does so is unclear. We used single-molecule multiwavelength fluorescence microscopy to test how the diffusion-limited binding kinetics of core RNAP to non-promoter DNA differ from those of mutant RNAPs that lack one or both α-CTDs. We find that even though α-CTDs and their tethers are small compared to the complete RNAP molecule, the presence of two α-CTDs accelerates DNA binding by ∼10-fold and ∼55-fold respectively relative to RNAP constructs in which one or both α-CTDs are deleted. In contrast, the presence of α-CTDs did not have a detectable effect on RNAP-DNA complex lifetimes in the absence of RNA synthesis. We explain how α-CTDs achieve the dramatic acceleration of RNAP binding to DNA using a quantitative three-state kinetic model that includes a transient binding intermediate where only the α-CTD(s) are bound to DNA, tethering the rest of the RNAP in the vicinity of DNA. The model and assumed parameters are validated using Brownian dynamics simulations of the DNA association reactions for two-, one-, or zero-CTD RNAP constructs. The combination of single-molecule experiments, mathematical theory, and simulations suggests that adding a flexible DNA-binding tether is a general physical mechanism which can accelerate the diffusion-limited binding of a large protein like RNAP to DNA and quantitatively defines the conditions under which this acceleration can occur.

**Significance Statement:** Large enzymes that must associate with DNA to perform their biological functions are expected to bind DNA only slowly because of slow enzyme diffusion. However, some DNA-binding enzymes have one or more additional small DNA-binding domains that can diffuse rapidly but are attached to the enzyme through an unstructured flexible tether. Combining single-molecule experiments, theory, and computation, we present evidence for a general mechanism by which such tethered domains can accelerate enzyme binding to DNA by orders of magnitude, while not significantly changing the duration of the DNA-bound state. We demonstrate this effect for the flexibly-tethered α-subunit C-terminal domain of bacterial RNA polymerase. The mechanism may allow the polymerase to rapidly bind DNA while minimizing non-functional sequestration on the genome.

## Introduction

In a variety of biological processes, large proteins or protein complexes must rapidly bind to DNA. The rates of such binding reactions is fundamentally limited by the protein-DNA collision frequency, but binding typically occurs in only a small fraction of the collisions in which the orientation of the DNA-binding site on the protein is appropriate to allow productive binding (1, 2). Mechanisms that increase the fraction of collisions that are productive, such as electrostatic steering [e.g., (3)], facilitated diffusion (4), or binding by unstructured protein domains, can potentially increase the overall binding rate, but the effects of these improvements are sometimes small (i.e., less than ∼2-fold), particularly at physiological ionic conditions (2, 5, 6).

In contrast to protein-DNA complexes with an interface formed from an unstructured domain, some proteins have small, globular DNA binding domains tethered to the bulk of the protein by an unstructured, flexible linker. The RNA polymerase enzyme from a broad range of [but not all (7)] bacteria species contains two such structures, the C-terminal domains of the α subunits (α-CTDs) (8, 9). In the RNAP holoenzyme which contains the sigma promoter recognition subunit, the α-CTDs and linkers have sometimes been modeled in complexes in which they are bound to DNA (10), but there are no direct observations of their structures in the absence of DNA. However, structural predictions (Fig. 1) reveal the α-CTDs (Fig. 1A, magenta), a ∼12-amino acid flexible linker (Fig 1A, yellow; Fig. 1B), and the N-terminal domains of the α subunits (Fig. 1A, cyan/blue). The α-CTDs are not required for the basic RNAP functions of transcription initiation and elongation. However, the α-CTDs are required for the function of some transcription activator proteins (11, 12), and they also can accelerate initiation of transcription, particularly at promoters that contain an AT-rich upstream promoter (UP) element (13–15).

**Figure 1:**
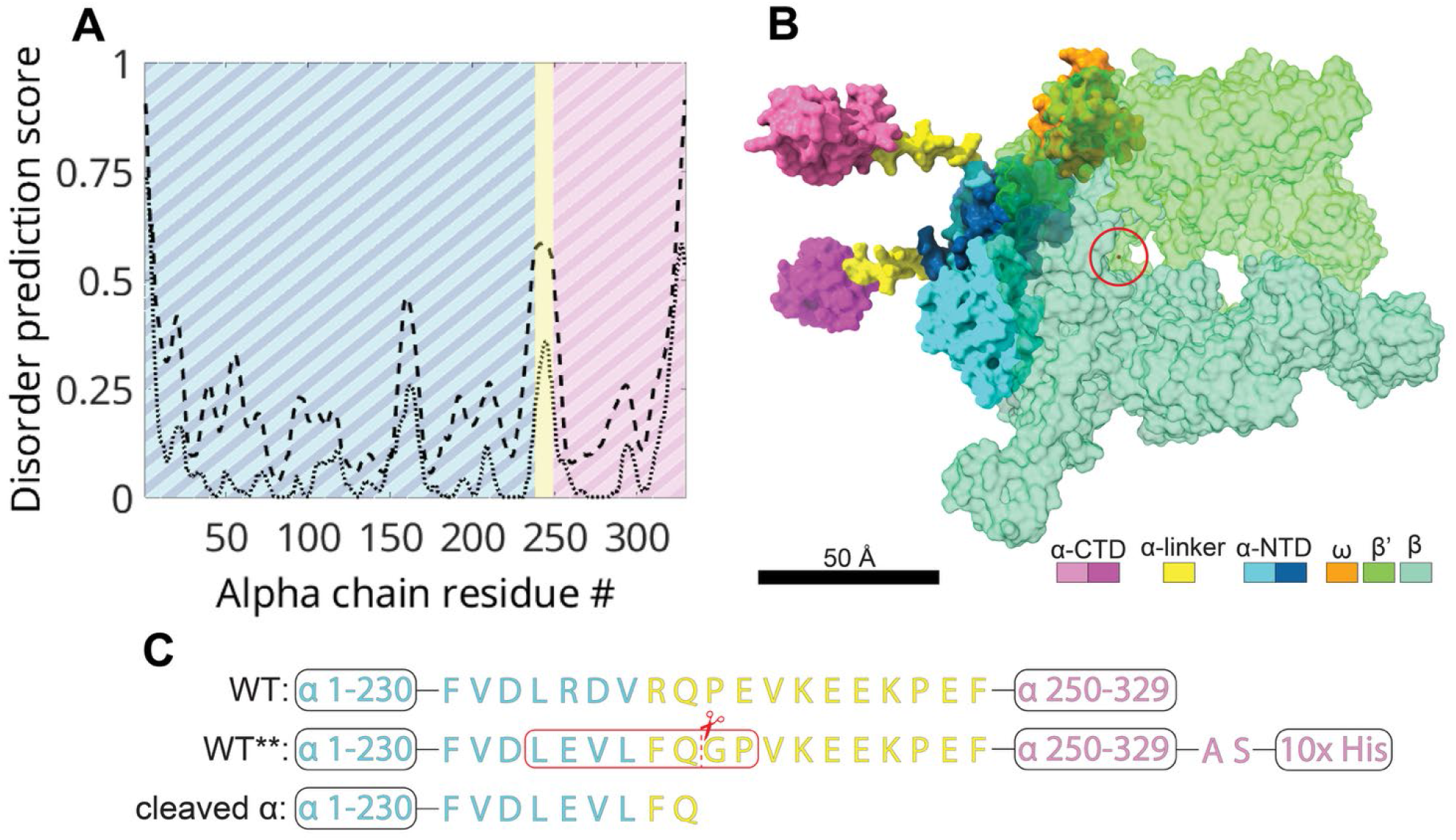
Predicted structure of core RNAP in the absence of DNA showing the complete α-subunits including α-CTDs and α-linkers. **(A)** Predictions of the intrinsically disordered segments of the α subunit sequence. Disorder scores are from PrDOS (40) (dashed line) and MetapredictV2 (41, 42) (dotted line). Colors indicate the extents of the α-NTD (cyan/blue), α-linker (yellow; 12 amino acids), and α-CTD (magenta) inferred from the scores (see Methods). The linker residue identification agrees closely with those from NMR studies on the isolated α subunit (43). **(B)** Experimentally determined 3-D structure of the core *E. coli* RNA polymerase β, β′, and ω subunits (PDB 7MKP) together with the docked structures of the two α subunits predicted by Alphafold 3 (44). The RNAP active site is circled. **(C)** Amino acid sequences of WT, WT**, and protease-cleaved α subunits, color-coded to match (A) and (B). The protease recognition sequence (red box) and cleavage site (dashed line) are marked.

α-CTDs are thought to accelerate early steps in transcription initiation, and single-molecule results suggest that an effect of α-CTD is to accelerate initial binding of RNAP to UP element-dependent promoters (13, 15). However, mechanisms of α-CTD involvement in transcription initiation are complex, and have been shown in various contexts to involve sequence-dependent interactions with UP elements, sequence-independent interactions with other DNA segments, interactions with σ^70^, and interactions with activator proteins (14–16).

To simplify analysis of the role of α-CTD in RNAP-DNA binding, we examined the formation of an initial RNAP-DNA complex, the post-termination complex (PTC). In the canonical transcription cycle, the PTC is formed after intrinsic termination concomitant with release of the RNA transcript (17–20). The PTC can also be formed directly from promoter-independent binding of core RNAP to DNA. The PTC is a long-lived, stable complex (18–20) that can adopt structures with the DNA positioned in RNAP similar to the position of DNA in the closed-or open-promoter complexes (21), but its formation does not require nucleoside triphosphates, a sigma factor, or a promoter sequence. RNAP can randomly slide long distances on DNA in the PTC state, indicating that its interaction with the DNA is largely sequence-independent.

Here, we used single molecule methods to directly compare the kinetics of DNA binding and dissociation by wild-type RNAP (containing two α-CTDs) and mutants containing one or no α-CTDs. Starting with the no-CTD construct, the addition of one or two α-CTDs progressively speeds up RNAP binding to DNA but does not affect the observed durations of RNAP dwells on the DNA. Kinetic and Brownian dynamics modelling show that these data are consistent with the formation of a short-lived intermediate in which the α-CTD is bound to the DNA and tethers the non-CTD portion of RNAP near the DNA. The formation of the intermediate is followed by the rapid full association of the tethered RNAP with the DNA. The combined single-molecule experiments and theory show that the presence of small, tethered DNA binding domains is a general mechanism that can be used to accelerate diffusion-limited binding of large proteins to DNA.

## Results

### Experimental determination of the effects of RNAP α-CTD on DNA association kinetics

To use single-molecule fluorescence microscopy to examine the effects of the α-CTD on DNA-binding properties we prepared (Figure S1; see Methods) *Escherichia coli* RNAP constructs each with a fluorescently-labeled SNAP tag fused to the C-terminus of the β′ subunit. In addition to the wild type (WT) construct (which has two α-CTDs; Fig. 1B) we also prepared a construct (WT*) in which both α subunits had a specific protease cleavage site substituted into the α-linker and a C-terminal His-tag (Fig. 1C) (22). Partial or complete protease digestion of WT* followed by immobilized metal affinity chromatography were then used to separate fluorescent RNAPs which have two α-CTDs (WT**), a CTD on only one of the α subunits (one-CTD), or that completely lacks CTDs (zero-CTD). All constructs were active in transcription (see Methods).

We then examined each RNAP construct in an experiment (Fig. 2A) in which we allowed the enzyme to bind to a surface-tethered, dye-labeled 586-bp circular DNA. At 1 nM WT protein in solution, we observed brief, reversible binding interactions at the locations of DNA molecules on the slide surface (e.g., Fig. 2B, Fig. 2C, Fig. S2). In the same recordings, we also monitored binding at randomly selected control locations that lacked labeled DNA molecules. Binding was more frequent at the DNA locations than at the control locations, indicating that some DNA-specific binding was observed at the former.

**Figure 2:**
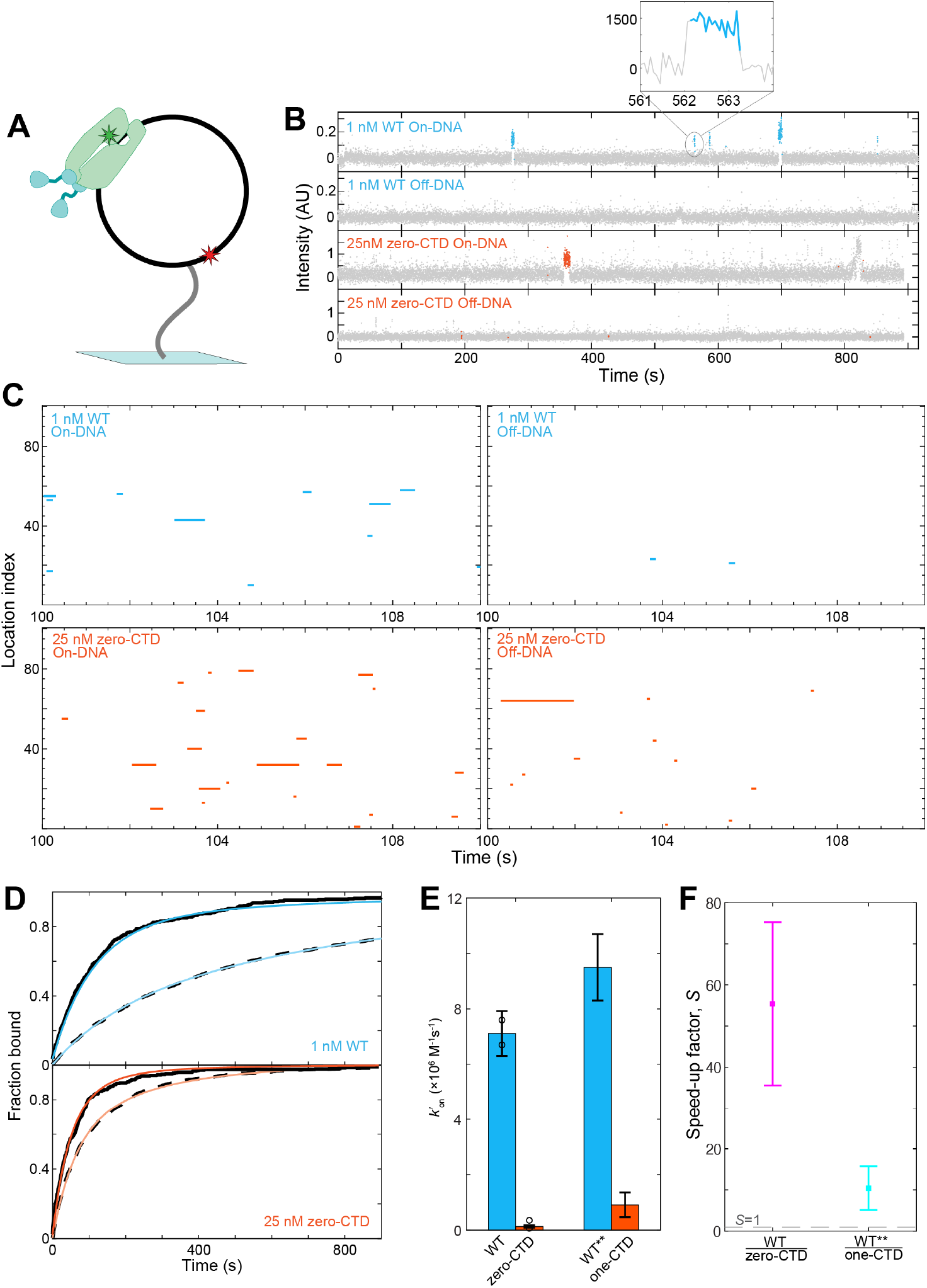
Single-molecule colocalization measurements of the association kinetics of RNAP constructs with DNA. **(A)** Cartoon of experiment design (not to scale). A 586-bp circular double-stranded DNA (black) labeled with a Cy5 is tethered to the surface of a microscope coverslip through interaction of a biotin modification of the DNA with a streptavidin-biotin-polyethylene glycol linkage (gray) to the slide surface. RNAP molecules (green) labeled with a green-excited dye are visualized binding to the surface-tethered DNAs by colocalization of the green and red fluorescence spots. **(B)** Example fluorescence intensity records recorded at the locations of a single DNA molecule (on-DNA) and at control locations lacking DNA molecule (off-DNA) from experiments with the indicated concentrations of WT and zero-CTD RNAP. Color and gray points represent times at which there is or is not a colocalized fluorescent RNAP spot **(C)** Rastergram plots of 100 randomly selected DNA or control locations from the experiments in (B). Color indicates presence of a co-localized RNAP spot. Each plot shows a magnified view of a portion of the time records in order to display detail; high resolution views of the complete time records are provided as supplemental data files. **(D)** Cumulative distributions of the fraction of DNA (solid black line) or control locations (dashed line) that exhibited binding of an RNAP molecule prior to the indicated time in the experiment (black) and fits (color) to a model that assumes exponential DNA-specific binding (see Methods). Number of observations and fit parameters are reported in Table S1. Experiments on one-CTD analogous to those in (B-D) are shown in Fig. S2. **(E)** Second-order association rate constants (± S.E.) for binding to DNA by zero-CTD RNAP, one-CTD RNAP, and RNAP constructs (WT, WT**) with two CTDs. Where points are shown, they are values from separate experiments, the bar is the weighted mean, and the error bar is the S.E. of the weighted mean. Data (see Table S1) are DNA-specific binding rate constants from the fits in (D), Figure S2B, and Figure S2C. **(F)** Data from (E) plotted as the Speed-up factor (± S.E.), the ratio of the association rate constants when both CTDs are present vs. when CTDs are reduced to zero or one. A value of *S* = 1 (dashed line) would indicate no speed up.

In initial experiments with the zero-CTD protein, we observed less binding than with WT. Therefore, we did experiments at higher concentrations of zero-CTD and one-CTD proteins (25 and 2.65 nM, respectively) than those used with the WT. The high protein concentrations resulted in more binding at off-DNA control locations (Fig. 2C, Fig. S2A). However, as for the WT we observed more binding at on-DNA than off-DNA locations, indicating that DNA-specific binding was seen. To quantitatively compare the association rate constants for the three proteins binding to DNA, we measured the time in the experiment at which each DNA molecule first bound RNAP and fit this data to a first-order binding model (Fig. 2D, Fig. S2B, Table S1). The fits demonstrate that the second order rate constant *k’_on_* (Fig. 2E) of WT RNAP was increased by more than 10-fold over one-CTD and by roughly 55-fold over zero-CTD (Fig. 2F). This change cannot be explained by a change in the linear diffusion coefficients of the molecules, which the presence of the α-CTDs is expected to decrease, not increase, and then only by a small (<< 2-fold) amount. The change also cannot be explained by removal of the α subunit C-terminal His tags, since the *k’_on_* value for WT (which lacks the tags) is not appreciably different than that of WT**, in which the tags are present (Fig. 1C). Thus, the presence of an α-CTD results in an order of magnitude increase in association kinetics, and two α-CTDs are substantially more effective than one.

To further investigate effect of the α-CTDs on RNAP-DNA interaction kinetics, we also measured how long the bound RNAP molecules remained on DNA before dissociating. Tenenbaum et al. (17) reported multi-exponential dwells of core RNAP on DNA, but did not fully define the dwell time distribution because of the limited dynamic range of individual dwell time experiments. Here, we measured dwells at both high (15 frames per second) and low (1 s duration frame every 5 s) time resolutions to increase the dynamic range of the measurements (23), and we performed the measurements on both WT and zero-CTD RNAPs (Fig. 3). The semilogarithmic dwell time distribution plots showed roughly linear regions in time ranges >∼100 s (Fig. 3, blue) and between ∼1 and ∼100 s (white), similar to the two-component distribution seen previously (17). The ∼100 s dwells were not seen in the high time resolution data because they are shortened by photobleaching. Conversely, we also observed an additional dwell time component with typical dwells < ∼1 s (Fig. 3, inset), which could be detected only at high time resolution. The multiple components indicate the presence of multiple kinds of RNAP-DNA complexes, consistent with the observation that core RNAP can stably bind to fully duplex DNA but also can open unwound DNA bubbles of at least two different sizes (21). Importantly, the WT and zero-CTD RNAP have essentially indistinguishable dwell time distributions. This suggests that WT and zero-CTD RNAP form similar complexes on DNA and that α-CTD-DNA interactions do not significantly increase the kinetic stability of any of the RNAP-DNA complexes that can be detected in these experiments.

**Figure 3:**
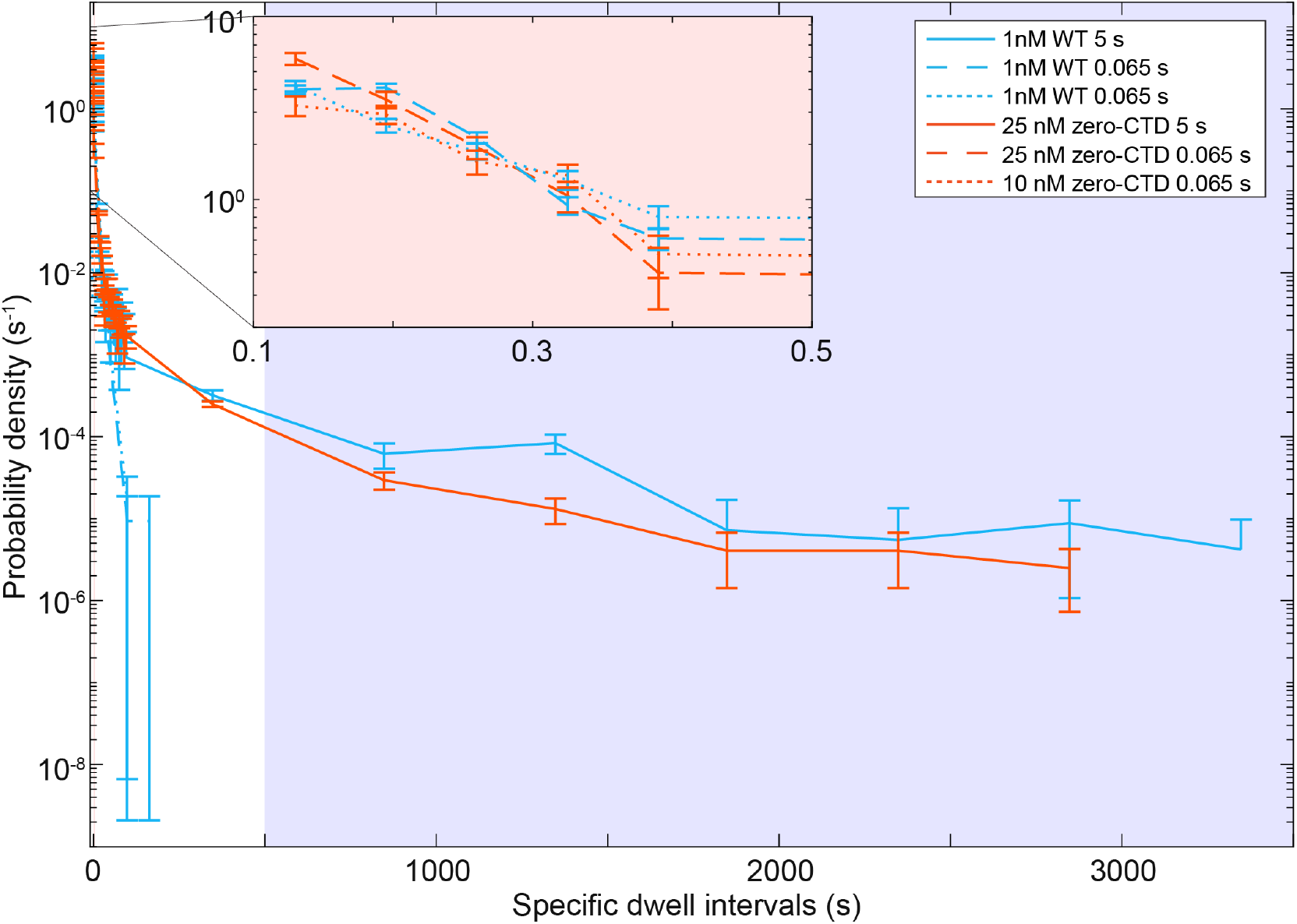
Distributions of WT and zero-CTD RNAP dwell times on DNA. Dwell times were measured in separate experiments at low (5 s) and high (65 ms) time resolutions; plots show the DNA-specific probability densities ± S.E. Inset: magnified view. Background colors highlight three widely different timescales on which dwells are observed. Only dwells that lasted for at least two consecutive frames were included.

### Kinetic model for binding acceleration by a flexible tether

The most striking result of the single-molecule experiments is that the presence of both or even one α-CTD dramatically accelerates RNAP binding to DNA (Fig. 2F). To explain this result, we propose a simple kinetic scheme for binding of WT or one-CTD RNAP to DNA (Fig. 4A, top) and examine whether it can feasibly explain the data. In the scheme, we consider a simplified model of RNAP as consisting of two domains, the transiently and weakly binding α-CTD and the rest of the RNAP that binds more strongly. The two domains are connected by a flexible polymer tether. Our scheme postulates a transient intermediate in which only the CTD is bound. For the WT enzyme, we simplify the model by treating the two CTDs as a single binding domain. For the zero-CTD enzyme, we treat the binding as a simple one-step process (Figure 4A, bottom).

**Figure 4:**
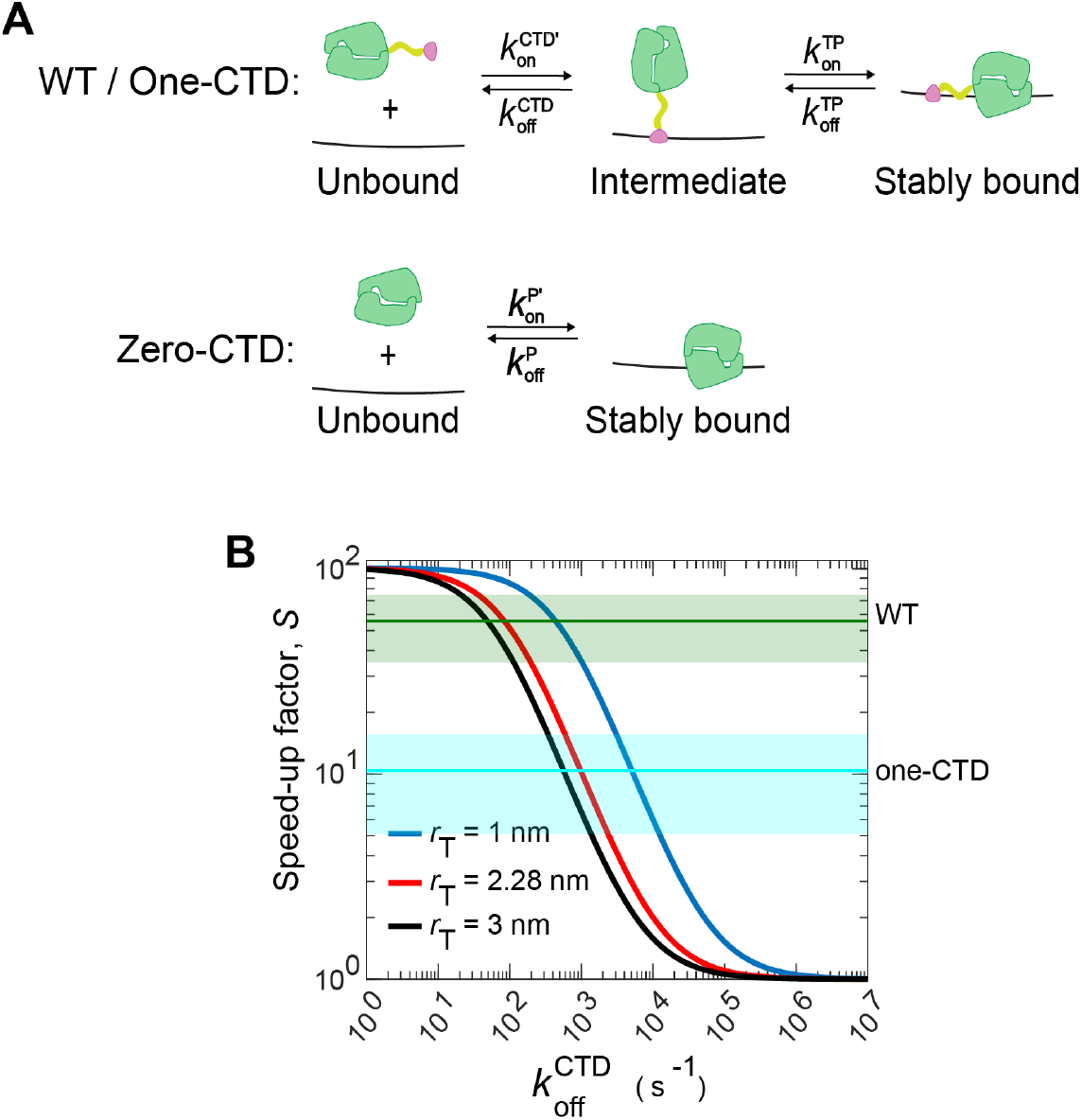
Kinetic modeling of RNAP-DNA binding. **A)** Kinetic models for tether-mediated binding of WT or one-CTD (top) and for conventional binding of zero-CTD RNAP (bottom) with DNA. Primes denote second-order rate constants; all other rate constants are first-order. Cartoons depict the CTD (purple), flexible linkers (yellow), and the tethered part of RNAP (green). **(B)** Speed-Up Factor (*S*) as a function of the α-CTD dissociation rate constant computed using equation 4. The exact length of the linker is not known, but it is estimated to be between 1 nm (solid blue line) and 3 nm (solid black line). The speed-ups relative to the zero-CTD are shown as horizontal lines: green, WT; light blue, one-CTD.

The lifetime data (Fig. 3) shows similar dwell time distributions for WT and zero-CTD, so we assume that the intermediate state has a lifetime << 65 ms, i.e., shorter than the time resolution of the experiment and the lifetime of the stably bound species. We also assume that the apparent first-order rate constant for binding the tethered part of RNAP to the DNA from the intermediate state is much larger than the rate constant for binding of the zero-CTD RNAP from solution. This is a consequence of the effective local concentration of the tethered part of RNAP around its tether site on the DNA being much larger than the typical (micromolar) concentration of polymerase in solution. For example, assuming that the α-CTD linker is a freely jointed polymer chain made of *N* = 12 amino acids (Fig. 1A), we can estimate its end-to-end distance as 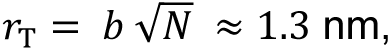 where *b* ≈ 0.38 nm is the distance between two adjacent amino acids in a peptide chain (24). However, this approach underestimates the end-to-end distance, as it assumes that the bond angles between adjacent amino acids are unconstrained. Using a machine-learning based predictor (25) for the conformations of an intrinsically disordered protein region we obtain an end-to-end distance *r*_T_ ≈ 2.3 nm. To accommodate the uncertainties in our determination of *r*_T_, for our estimate of the speed up factor in Figure 4B, we use a range of values: 1 nm ≤ *r*_T_ ≤ 3 nm. This yields an estimated effective local concentration of the tethered part of RNAP 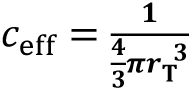 between 10 – 400 mM.

Based on the estimated *r*_T_ for the α-CTD linker, we can compute the apparent first order rate constant for association of the tethered part of RNAP to a site on the DNA as

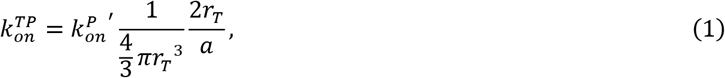

where *k^p’^_on_* is the second order rate constant per base pair of DNA for stable binding of the zero-CTD RNAP. A simple estimate of this rate is given by the Smoluchowski formula *k^p’^_on_ =4πDa* where *D* is the diffusion constant of the polymerase and *a* ≈ 0.33 nm is the size of a binding site on the DNA. In equation 1 we also consider the number of accessible binding sites for the tethered part of the RNAP, 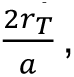 where 2*r_T_* is the length of the DNA accessible to the tethered polymerase.

Given the kinetic scheme in Figure 4A, we can compute the effective second order rate constant per base pair for the formation of the stably bound WT RNAP as:

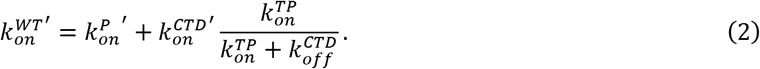

This formula assumes that the binding rate from the unbound to the intermediate state is much slower than the binding rate from the intermediate state to the stably bound state, which is a consequence of the much higher local concentration of tethered part of RNAP around the DNA in the intermediate state, as discussed previously. In addition to the binding from the pathway in Fig. 4A (top) we include the direct binding pathway with rate constant *k^p’^_on_*, analogous to the pathway shown in Fig. 4A (bottom).

With this formula in hand, we can express the speed up factor, i.e., the ratio of the effective association rate constant for the formation of the stably bound state of the WT polymerase (*k*^WT*’*^*_on_*) to the association rate constant for the formation of the stably bound state of mutant RNAP (*k^p’^_on_*) as:

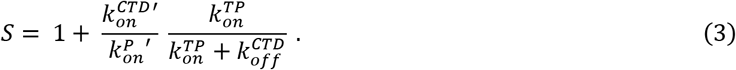

To make specific numerical predictions of *S* based on this formula, we assume that when α-CTD alone binds to DNA, it imposes little constraint on the orientation of the tethered part of RNAP, whereas in the stably bound species, RNAP makes multiple contacts with the DNA that constrain its orientation. An analogous assumption was made by Northrup and Erickson for protein-protein binding (26). Based on that work, we estimate the ratio of the on-rate constants 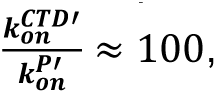 an assumption we validated by performing Brownian dynamics simulations (see below). With this estimate in hand and based on the measured speed-up ratio from zero-CTD to two-CTD RNAP, *S* = 55 ± 20, we calculate the partition ratio from equation 3 to be 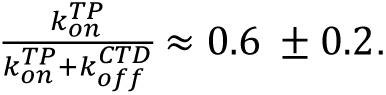.

Substituting equation 1 fo *k^TP^*_on_ into equation 3, the formula for *S* becomes

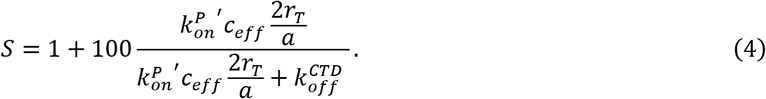

For the rate parameter *k^p’^_on_* we use the value 3.8 × 10^2^ M^-1^ s^-1^ bp^-1^ which we obtain by taking the measured second-order association rate constant (2.0 × 10^5^ M^-1^ s^-1^; Figure 2E) for the zero-CTD polymerase and dividing it by the length of the DNA used in our single molecule experiments (586 bp).

We plot equation 4 in Figure 4B as a function of the off rate for the CTD (*k*^CTD^_off_) for different values of the tether length, *r*_T_. In this figure we indicate the measured *S* (Figure 2F) from our single-molecule experiments by the two horizontal lines, one corresponding to the one-CTD RNAP (*S* ≈ 10) and the other to the WT RNAP (*S* ≈ 55); the speed-up for the zero-CTD is *S =* 1 by definition.

This comparison of the model to the data from the single-molecule experiments suggests that the dissociation rate for the intermediate state when two CTDs are present is *k^CTD^_off_* ≈ 40-400*s*^-1^, obtained by reading off the intersection of the WT (green) line with the theoretical curves (Figure 4B). For the one-CTD RNAP, the dissociation rate of the intermediate state is similarly predicted to be about an order of magnitude greater. This is expected due to the anticipated larger stability of the intermediate state in Figure 4A when two CTDs are bound compared to the state with only one CTD bound. The predicted *k*^CTD^_off_ values correspond to dwell times in the intermediate state that are 25 ms or shorter, which is below the time resolution of our experiment (65 ms). This rationalizes the failure to observe dwell times that correspond to the intermediate state in Figure 3.

### Brownian dynamics of accelerated binding by a flexible tether

The previous analysis shows that the kinetic model for tether-mediated binding can explain the experimental data, but this conclusion is based on assumptions, such as treating the effect of the CTD tether as imposing an effective concentration, and the assumed 100-fold larger rate of the CTD association with DNA compared to the rate of association of the non-CTD part of the polymerase. To test these assumptions, we performed Brownian dynamics simulations of a coarse-grained model of the RNA polymerase and the DNA to which it binds.

In the simulation, we represent the non-CTD portion of the RNAP as a triangle whose vertices serve as contacts to the periodic array of cognate binding sites on the DNA (Fig. 5A). We chose to use a triangle since three points uniquely define the position and orientation of the RNAP relative to the DNA. Also, the configuration when all three vertices of the triangle are in contact with the binding site on the DNA represents the stably bound state of the RNAP and DNA. The α-CTD is represented as a point at the end of a polymer tether. The polymer tether, whose root-mean-square end-to-end distance is *r_T_*, connects the triangle to the point representing the α-CTD. Interactions between each of the four contact points representing the RNAP and their DNA targets are modeled as a short-range potential. The motion of all four points representing the RNAP is diffusive (see Methods for details). RNAP is confined within a sphere with reflecting boundary conditions, with the size of the sphere setting the RNAP concentration in solution.

**Figure 5:**
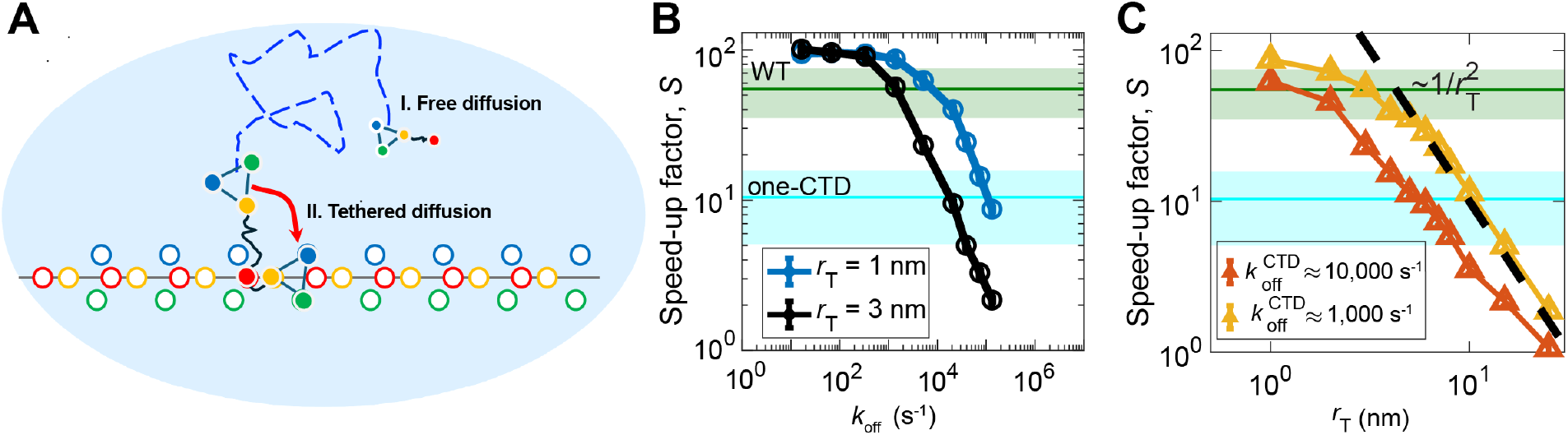
Brownian dynamics simulation of RNAP-DNA binding. **(A)** Schematic of the Brownian dynamics model of RNAP association with DNA. The solid green, yellow and blue circles represent the non-CTD part of the polymerase, while the red circle, attached via the black tether, is the α-CTD. The open circles represent the corresponding binding sites on the DNA. The association reaction consists of two steps: (1) RNAP first performs 3-D diffusion (dashed line) until the CTD attaches to its target (red open circle), and (2) once the CTD is bound, the non-CTD part of RNAP can then quickly attach to the target by performing tethered diffusion (red arrow). **(B)** Speed-up factor *S*, the ratio of the average time to first binding without and with the CTD tether, for different tether lengths. The horizontal lines show the speed-up factors obtained from the single-molecule experiments. **(C)** Speed-up factor as a function of the tether length for two different values of *k*^CTD^_off_. The dashed black line corresponds to the scaling relation *S* ∼ *r_T_*^−2^ (see text). Simulations are described in the Methods section.

We ran the simulations with and without the α-CTD for different values of *r_T_* and *k*^CTD^_off_. To vary *k*^CTD^_off_, we varied the binding energy of the point representing the α-CTD with its cognate site on the DNA. For each binding energy, *k^CTD^_off_* was computed from the measured mean escape time of the RNAP from the state in which only the CTD is bound.

For each set of conditions, we estimated the mean time to form the stably bound RNAP-DNA complex by averaging 200 individual RNAP-search simulations, each starting with the RNAP at a random position within the confinement sphere. Each simulation continued until all three contacts representing the non-CTD portion of RNAP reached their target. The speed-up factor (Fig. 5 B and C) was then computed as the ratio of the mean times to form the complex without and with the CTD.

The results of the Brownian dynamics are consistent with our kinetic model. First, we measured the distributions of times to go from the unbound to intermediate state, from the intermediate to the stably bound state, and from the intermediate to the unbound state. The distributions are approximately exponential (Fig. S4) which justifies the use of the simple Markov state model (Fig. 4A) in the kinetic analysis. Second, we find a similar relationship between the speed-up factor and k_off_^CTD^ in the Brownian dynamics simulation as we did in the kinetic model (compare Fig. 5B and Fig. 4B). As in the kinetic model, the speed-up ratios measured in experiments again imply a short-lived intermediate with only CTD bound. The Brownian dynamics predict an intermediate lifetime less than a millisecond (see the intersection of simulation curves with experimental results for WT and one-CTD RNAPs in Fig. 5B). As was inferred from the kinetic model, this is well below the time resolution of the experiment, and consistent with the failure to observe the difference in RNAP dwell times for WT and mutant polymerases. The Brownian dynamics also confirms the prediction that the scaling of the speed up with the length of the CTD tether, *S* ∼ *r_T_*^−2^ (Figure 5C). This scaling regime is obtained from equation 4 in the limit when 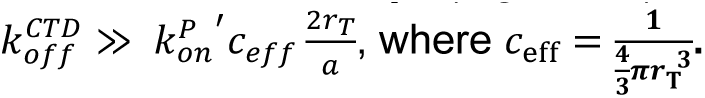 Thus, the simulations support the assumptions we made in deriving equation 4.

Brownian dynamics simulations also allowed us to compare binding dynamics for one and two-CTD RNAPs. Consistent with experiments, the simulations show that the speed-up factor is greater for the two-CTD RNAP (WT) than the one-CTD case, regardless of the length of the flexible linker (see Supplementary Text). Also, as assumed in our kinetic model, the different effects of having one vs two CTDs on the speed up can be captured by a change in the off rate for CTD unbinding from the DNA. This further justifies the use of the simpler, one-CTD RNAP in the theory leading up to equation 4.

## Discussion

The experiment and theory results presented here show that the α-CTDs speed up the diffusion-limited binding reaction of RNAP to DNA and can explain the physical mechanism by which this acceleration is achieved. In general, we would expect that adding protein domains to the non-CTD portion of RNAP would slow its translational and rotational diffusion. This would slow down, rather than speed up, the diffusion-limited association reaction. Instead, our results instead imply that the slowing effects of the added α-CTDs are greatly overcome by the effect of the α-CTDs in increasing the fraction of collisions between RNAP and DNA that produce stable binding. The kinetic and Brownian dynamics models together show that two features of the tethered α-CTDs explain this increase. First, once RNAP is within a short distance of the DNA, the α-CTDs are able to bind DNA faster than the non-CTD portion of RNAP. This is expected both because the α-CTD translational and rotational diffusion are faster and also because its DNA binding interface is a larger fraction of its surface area than it is for the non-CTD portion of RNAP. Second, once the non-CTD portion of RNAP is tethered by the α-CTD to the DNA, the rate of its binding attempts to form the stably bound state increases. Importantly, the models explain the effect of the α-CTDs purely in terms of diffusion dynamics, without reference to electrostatic attraction or steering. Thus, they are plausible descriptions of molecular behavior at physiological ionic conditions, in which long-range electrostatic forces are weak.

Our models suggest that during PTC formation, RNAP can bind to the DNA via the α-CTDs, first making a transient intermediate that cannot form in the absence of α-CTDs. An intriguing result of our experiments is that despite the presence of this intermediate, we do not observe any significant difference in the dwell times on DNA of core RNAPs with and without α-CTDs. This observation leads to two conclusions about the intermediate. First, it must be short-lived. Since we do not see an additional short component of the dwell time distribution due to the presence of α-CTDs (Figure 3), the intermediate lifetime must be significantly shorter than the 65 ms resolution of our fastest dwell time measurements. Second, the failure of the CTDs to affect the measured dwell time distribution implies that the two-CTD RNAP usually does not enter the stable binding state more than once before dissociating from the DNA. Multiple entries into the stably bound state would increase the average dwell time over that seen with zero CTDs (Figure 4A, bottom).

We estimate that our data (Figure 3) rules out a two-CTD induced increase in the short component dwell time of more than a factor of two. This implies that the partition ratio for the intermediate, 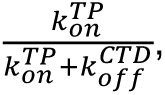 is less than ∼ 0.5, consistent with the value of 0.6 ± 0.2 calculated earlier. Thus, the properties of the α-CTD seem optimized to promote rapid binding of RNAP to DNA (which requires the partition ratio to be relatively large), while minimizing the dwell time of non-specifically bound RNAP (which requires the partition ratio to be relatively small). Both of these effects may be adaptive for rapid bacterial growth in nutrient-rich environments.

α-CTDs function at multiple different stages of the bacterial transcription cycle. In the elongation complex, the α-CTDs do not appear to interact with DNA but instead contact the NusA elongation factor (27). In transcription initiation, α-CTDs interact with upstream DNA (either with or without a UP element) and also can contact activators and region 4 of σ^70^ (14, 15, 28). α-CTDs are also involved in the post termination complex, where their interactions with DNA are thought to promote the flipping process that leads to reinitiation of transcription on oppositely oriented promoters (18, 29). The stimulation of transcription by α-CTDs is consistent with the acceleration of RNAP-DNA binding demonstrated here, but α-CTDs may also promote additional steps in the transcription cycle.

In addition to showing how the α-CTDs can accelerate binding of core RNAP to DNA, the theory and computational work presented here suggest that having a small, tethered DNA binding domain could serve as a general mechanism for acceleration of the association of a larger DNA-processing protein with DNA. Searches of structural databases reveal multiple candidate proteins that are annotated as having have small DNA-binding linked to larger DNA binding domains by intrinsically disordered linkers (Table S3). It remains to be seen in these cases whether the presence of the small domain serves to dramatically accelerate binding of the large domain, as we observe for the effect of the α-CTD on RNAP binding to DNA.

## Methods

### Proteases

Human rhinovirus 3C protease fused to a GST tag (ppx) (30) was a kind gift from Bruce Goode’s lab. The 3C protease fused to a decahistidine tag (his-ppx) was expressed from plasmid His-ppx3C (kind gift from Seth Darst’s and Elizabeth Campbell’s labs) in *Escherichia coli* BL21 (DE3) by growing for ∼4 hr at 37°C to OD_600_ ∼0.6 then reducing the temperature to 20°C and inducing with 0.5 mM IPTG for ∼16 hr. The cells were then spun at 4,500 × *g* for 20 min at 4°C then resuspended in 1 ml per g cells lysis buffer (20 mM Tris, 5% glycerol, 0.5 M NaCl and 1 mM 2-mercaptoethanol). Cells were then lysed by sonication and DNAse I (0.5 mg/ml) and RNAse A (1 mg/ml) were introduced and the lysate incubated for 30 min on ice. The solution was then spun at 267,000 × *g* for 20 min. The supernatant was then loaded into a 1 ml HisTrap column (Cytiva Life Sciences). The column was washed with wash buffer (20 mM Tris-Cl pH 8.0, 1M NaCl, 5% [v/v] Glycerol, 1 mM 2-mercaptoethanol) containing 20 mM imidazole and eluted with a gradient from 20 mM to 500 mM imidazole in wash buffer. The fractions were pooled and stored in 25% glycerol at-80° C.

### Dye-labeled β’-SNAP RNAP (WT RNAP)

*E. coli* core RNAP (αββ’ω) with consecutive SNAP and hexahistidine tags on the C-terminus of β’ was expressed (31) and purified (32) to give RNAP-SNAP. RNAP-SNAP was labeled with SNAP-surface 549 dye (New England Biolabs) (18) to produce “WT” RNAP.

### RNAP with cleavable α-linkers and β’-SNAP (WT* RNAP)

Plasmids pEcRNAP5 (Addgene #128939), pACYCDuet-1_Ec_rpoZ (Addgene #128837), and E. coli strain BL21(DE3)T-X_234-241_H (22) were generously provided by Seth Darst’s lab. To make the WT* RNAP construct, which has α subunits with cleavable α-linkers and C-terminal decahistidine tags and a dye-labeled SNAP tag fused to the C-terminus of β’ (β’-SNAP), we first PCR amplified the SNAP sequence from vector pSNAP-tag (T7)-2 (New England Biolabs) using primers 5′-GGG CGG TTC TGA TAA CGA GCT CGA CAT GGA CAA AGA TTG CGA AAT GAA AC-3′ and 5′-TCA TAG CTG TTT CCT GTC ATC CCA GAC CCG GTT TAC C-3′. Next, the SNAP sequence was inserted into XhoI-cut pEcRNAP5 using the NEBuilder HiFi DNA Assembly (New England Biolabs) to make pECRNAP5-β’-SNAP. ECRNAP5-β’-SNAP and pACYCDuet-1_Ec_rpoZ were co-transformed into BL21(DE3)T-X_234-241_H.

To express and purify WT* RNAP, cells were grown in 6 liters of LB broth at 37°C to OD_600_ ∼0.6 then induced with 1 mM IPTG for 4 hours at 30°C. Next, the cells were spun down at 4,500 × *g* for 20 min at 4°C and resuspended in lysis buffer (50 mM Tris-Cl pH 8.0, 10 mM dithiothreitol (DTT), 5% [v/v] glycerol, 1 × Pierce protease inhibitor (ThermoFisher #A32955)) at 1 ml per gram of cells. The cells were lysed via sonication on ice for a total of 2 min. Afterwards, the cells were spun at 27,000 × *g* for 30 min at 4°C and the lysate was collected in a beaker on ice. 10 % polyethyleneimine (Polymin P) was slowly added to a final concentration of 0.6 % while being stirred at 4°C and was left stirring for an additional 25 min. The mixture was centrifuged at 27,000 × *g* for 1.5 hr at 4°C and the supernatant discarded. The pellet was then consecutively resuspended and centrifugated four times, twice in PEI wash buffer (50 mM Tris-Cl pH 8.0, 0.5 M NaCl, 10 mM DTT, 5% [v/v] Glycerol, 1 × Pierce protease inhibitor) then twice with PEI elution buffer (50 mM Tris-HCl pH 8.0, 1 M NaCl, 10 mM DTT, 5% [v/v] Glycerol, 1 × Pierce protease inhibitor). The supernatants from the two PEI elution buffer washes were pooled. Then ammonium sulfate to 0.35 g per ml PEI supernatant was slowly added with stirring at 4°C and left to stir for at least 12 hours. The mixture was then spun at 27,000 × *g* for 45 min at 4°C and the supernatant was discarded. The pellet was resuspended with 3 – 5 ml wash buffer, injected into a 1 ml HisTrap column (Cytiva Life Sciences), washed with wash buffer until the OD_280_ was stable, then eluted with 250 mM imidazole in wash buffer. The imidazole was then reduced 225-fold by cycles of concentration, dilution with wash buffer and re-concentration using an Amicon Ultra-15 centrifugal filter (Sigma-Aldrich). To the concentrated protein, glycerol was added to 30% [v/v] and DTT to 2 mM. The preparation was then flash frozen in liquid N_2_ and stored at-80°C.

### Dye-labeled RNAP with no α-CTDs (zero-CTD RNAP)

To make RNAP lacking α-CTDs, 10 μl his-ppx protease (∼40 μM) was mixed with 500 μl WT* (5 μM) and incubated overnight at 4°C to fully digest all α subunits. The solution was then passed through a 1 ml HisTrap column (Cytiva Life Sciences) with wash buffer (20 mM Tris-HCl pH 8.0, 1 M NaCl, 5% [v/v] Glycerol, 1mM BME) to remove the his-tagged α-CTD and his-ppx. To fluorescently label the protein, SNAP-Surface 549 (New England Biolabs) was added at a 1:4 protein-to-dye mole ratio and incubated for 30 min at room temperature. Unreacted dye was removed using a Centrispin 20 column (Princeton Separations) equilibrated in 20.7 mM Tris-Cl pH 8.0, 200 mM NaCl, 2 mM DTT. Glycerol was added to 30% [v/v] and bovine serum albumin (BSA) to 1 mg/ml. The sample was flash frozen in liquid N_2_ and stored at-80°C. WT and zero-CTD preparations were confirmed active in single-molecule transcription experiments (18) which showed that ∼50% and ∼37%, respectively, of the molecules in the preparations were active.

### Dye-labeled one-CTD mutant RNAP (one-CTD) and two-CTD RNAP control (WT** RNAP)

To prepare β’-SNAP RNAP with one CTD, WT* RNAP was diluted 1:5 with wash buffer (20 mM Tris-Cl pH 8.0, 1M NaCl, 5% [v/v] Glycerol, 1 mM 2-mercaptoethanol) supplemented with 5 mM DTT and then digested for 75 - 90 min with ppx protease at a 1:5 mole ratio of protease to RNAP. The sample was then loaded into a 1 ml HisTrap column (Cytiva Life Sciences), washed with wash buffer, and eluted with a 0 to 400 mM imidazole gradient in wash buffer at 0.3 ml/min, which took a total of 2 hours to complete. The different elution of proteins with one and two his-tags was used to purify RNAP species with one and two α-CTDs. The indicated column fractions (Figure S1) were depleted of imidazole as described for WT* RNAP and one-CTD RNAP. WT** RNAP, which has the same structure as WT* RNAP, was prepared by labeling, removing unreacted dye, and storing as described for zero-CTD RNAP. In the intact RNAP, the two α-CTDs are known to have some differences in structure and function (33), but for simplicity this distinction was ignored in preparation and analysis of the one-CTD mutant.

### DNA

To avoid the previously described tight binding of the RNAP to DNA ends (34), a non-supercoiled circular template DNA was made as described (35). The 586 bp DNA has no annotated promoters and contains two labels: Cy5 and biotin.

### Single-molecule observation of RNAP binding to DNA

Single molecule experiments were done on a micro-mirror total internal reflection (TIR) fluorescence microscope (36) at 532 nm, and 633 nm excitation wavelengths to observe labeled RNAPs, and DNA respectively. The microscope focus was automatically maintained (37). Observations were done in glass flow chambers of ∼15 μl volume passivated (36) with a mixture of mPEG-SG2000 and biotin-PEG-SVA5000 (Laysan Bio). All reagent additions and washes used 20 μl volumes. To facilitate later correction for stage drift, the chamber was first incubated with streptavidin-coated fluorescent beads (T-10711, Molecular Probes) in wash buffer (50 mM Tris-acetate, 100 mM potassium acetate, 8 mM magnesium acetate, 27 mM ammonium acetate, 0.1 mg ml^−1^ BSA (#126615 EMB Chemicals), pH 8.0) to a dilution of ∼1:400,000 before washing out the excess. Streptavidin (220 nM; #21125; Life Technologies) was then introduced in wash buffer, incubated 45 s, and the excess washed out. 190 pM labelled DNA was then incubated in the chamber for roughly 20 min before being washed out with wash buffer supplemented with an O_2_ scavenging system [4.5 mg/ml glucose, 40 units/ml glucose oxidase (Sigma Aldrich #9001-37-0), 1,500 units/ml catalase (Sigma Aldrich, #9001-05-2), and 2 mM DTT] and the microscope focus adjusted. Next, labelled RNAP (1 nM WT, 2.65 nM one-CTD, or either 10 or 25 nM zero-CTD) was introduced and the camera set to record. High time resolution image acquisition was done at 65 ms per frame where for ∼5 s the sample was excited with 633 nM to record the DNA locations and then for ∼15 min with 532 nm to observe RNAP binding. Laser powers (measured incident to the micro-mirror) were 10.8 mW and 7 mW for 532 nm and 633 nm, respectively. For low time resolution data, DNA locations were recorded for 5 s at 1 frame per second before recording the RNAP at 5 s intervals (20% duty cycle) for ∼1 hr with 300 μW micro-mirror incident laser power for both 532 nm and 633 nm.

### Measurement of RNAP-DNA association kinetics

The rate of association of RNAP with DNA (Fig. 2D-E, Table S1, Fig. S2) was measured using the general approach described (22) with some modifications. In summary, we first identified sites on the slide surface that had a fluorescent DNA molecule (on-DNA sites) and randomly selected negative control sites with no fluorescent DNA (off-DNA sites). At each site we measured the interval between the start of the recording and the time at which the first RNAP molecule was observed to bind. We assumed that both on-DNA and off-DNA sites have the potential to non-specifically bind RNAP to the slide surface; this non-specific binding was parameterized by maximum-likelihood fitting the off-DNA time-to-first-binding data to the probability density function

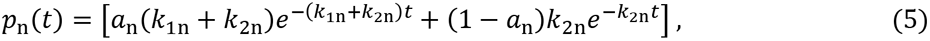

yielding parameter values of *a*_n_, *k*_1n_, and *k*_2n_.The on-DNA data from the same recording were then analyzed using a model that assumes that of the *N* on-DNA sites in the microscope field of view, only a fraction, termed the active fraction (*A*_f_), are capable of binding RNAP. The remaining fraction of DNA sites available to first bind RNAP at the start of the recording is then *A_f_* − *^nz^*/*_N_*, where *n*_z_ is the number of on-DNA sites already bound to RNAP. Using fixed values of a_n_, k_1n_ and k_2n_ determined from the above optimization of the off-DNA binding data (eqn. (5)) we determined the apparent first-order association rate constant *k*_on_ and the *A*_f_ by maximum-likelihood fitting the on-DNA time to first binding data using the probability density,

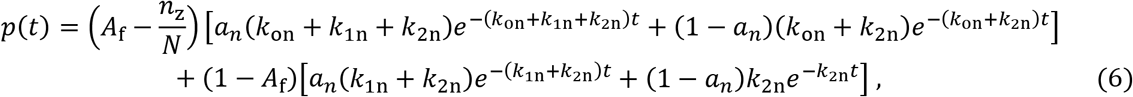

using a single global value of *A*_f_ for experiments done on the same slide with either WT and zero-CTD RNAP or WT** and one-CTD RNAP. Fit parameter standard errors were calculated by bootstrapping (5,000 samples).

### Measurement of RNAP-DNA dissociation kinetics

To calculate the DNA-specific dwell time distribution (Fig. 3), we corrected the measurements recorded for on-DNA sites using those from the off-DNA sites to remove the DNA-independent contributions to the binding at the on-DNA sites. The number of on-and off-DNA sites differed, so for each interval bin in the distribution we adjusted the number of off-DNA landings multiplying by the ratio *W*_s_/*W*_ns_, where *W*_s_ is the total wait time between on-DNA bindings at all sites (i.e., total interval when no RNAP is bound) and likewise *W*_ns_ is the total wait time between bindings at all off-DNA sites. Since the nonspecific surface binding also occurs at DNA sites, the number of DNA-specific bindings is (*n_s_* − then 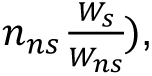 where *n*_s_ is the number of on-DNA bindings with durations contained in that bin and *n*_ns_ is the number of off-DNA bindings that occur within that same interval bin. Dividing this difference by both the bin width and the total number of DNA-specific bindings then yields the DNA-specific probability density for that bin as plotted in Fig. 3.

The uncertainty for the binding number associated with just the on-DNA bindings in a bin was computed using Δ*n_s_* = [*p* (1 − *p*) *N*]^0.5^, where *N* is the total number of on-DNA bindings, and *p* = *n_s_*/*N* is the probability that an on-DNA binding occurs with a dwell duration that places it within this bin under consideration. Similarly, the uncertainty in the off-DNA binding number is Δ*n_ns_* = [*p_ns_* (1 − *p_ns_*) *N_ns_*]^0.5^ where *N*_ns_ is the total number of off-DNA bindings and *p_ns_* = *n_ns_*/*N_ns_* is the probability that an off-DNA binding will occur with a dwell duration that places it within this same bin under consideration. The uncertainty in the adjusted number of off-DNA bindings is then Δ*n*_ns_ (*W*_s_/*W*_ns_). The uncertainty for the DNA-specific binding number for the bin will then be 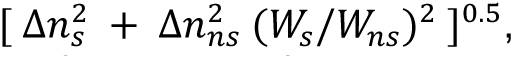 and dividing this number uncertainty by both the bin width and the total number of DNA-specific bindings yields the uncertainty in the DNA-specific probability density for that bin. That uncertainty in probability density computed for each bin then defines the sizes of the error bars appearing in the Fig. 3 distribution.

### Estimation of α-CTD linker length

To model the effects of α-CTD on RNAP-DNA association/dissociation kinetics, we first estimated the length of the α-CTD linker. The linker length was determined using AlphaFold and a number of different protein disorder predictors (Fig. 1B). The AlphaFold Protein Structure Database (38, 39) contains a structure for *E. coli* RNAP α subunit. The predicted local distance difference test (pLDDT) and predicted alignment error (PAE) were used to indirectly estimate the tether sequence. This led to a prediction of residues 235-250 for the tether. Since these predictions were indirect, the protein disorder predictors were used to verify these sequences. The protein disorder predictor PrDOS (40) predicted residues 236-249 to be disordered (disorder probability > 0.5). Metapredict V2 (41, 42) did not predict anything to be disordered in the region AlphaFold predicts the tether. However, around the region AlphaFold and PrDOS predict the tether, Metapredict V2 showed higher relative disorder. In this region, the maximum disorder score was 0.36. To determine residues in the tether, residues with a score of at least 0.18, half the maximum, were taken to be the tether. From this, Metapredict V2 predicted residues 238-249 for the tether. From these predictors and AlphaFold, residues 238-249 were taken to be the consensus tether. This assignment is consistent with the results obtained from NMR studies of the isolated α subunit and C-terminal fragments (43). To estimate the end-to-end distance of this sequence, the linker consensus residues were input to ALBATROSS (25), a machine learning model that predicts disordered protein physical properties such as radius of gyration, end-to-end distance, and asphericity. ALBATROSS estimated the linker to be 2.28 nm long. Brownian dynamics calculations used this length as the value for the tether length and also used values of 1 nm and 3 nm to account for uncertainties in the estimate.

### Brownian dynamics

The Brownian dynamics simulations were performed for the model shown in Fig. 5A, for which the confining sphere has a diameter of 500 nm. The rest length between the points on the equilateral triangle representing the non-CTD part of the RNAP is *l*_0_ = 5 nm, while the root-mean-square length of the polymer tether representing the CTD linker (*r*_T_) is varied. The RNAP binding sites on DNA are represented as a periodic array of 20 repeats of four points, each representing a binding site matched to the corresponding point on the RNAP. The distances between the points on the DNA (open circles in Fig. 5A) are matched to the rest spacings between the points representing the RNAP (filled circles in Fig. 5A). Details of this geometry are described in Fig. S3 and the parameters used in the simulations are collected in Table S2.

The over-damped Brownian dynamics for the displacement 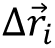 of the four points representing the RNAP, for a time step Δ*t*, is computed as:

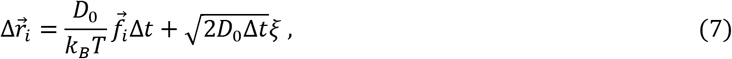

where *i* = 1, 2, 3 are the points on the triangle representing the non-CTD part of the polymerase (green, yellow, and blue filled circles in Fig. 5A) and *i* = 4 is the point that represents the CTD (red filled circle in Fig. 5A). *D*_0_ is the diffusion coefficient for each point and *ξ* is a random number drawn from the standard normal distribution. For simplicity, we work in units of time and energy for which *D*_0_ = 1 and *k_B_T* = 1; all lengths are measured in units of nm.

The force on each of the four points representing the RNAP is computed as 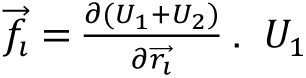 is the energy of interaction between the points on the RNAP and their cognate sites on the DNA, which is given by

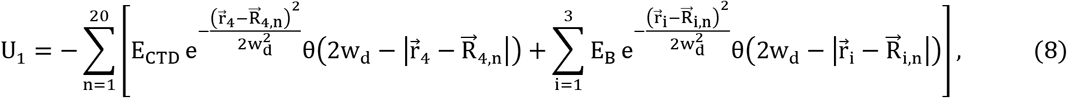

where *w_d_* = 0.5 nm is the target size. *θ* is the Heaviside function; if the distance between the point representing the RNAP and its cognate site on the DNA 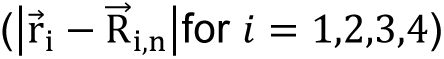 is larger than 2*w_d_*, the interaction energy is zero.

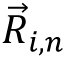 represent the positions of the binding sites on the DNA with a period of 5 nm, matching the rest length of the contacts, *l*_0_. The distance between the binding site for the CTD 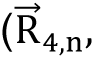 red open circle in Fig. 5A) and the closest site (yellow open circle in Fig. 5A) for binding the non-CTD part of the RNAP is taken to be 0.3 nm (Fig. S3).

The interaction energy between the non-CTD portion of the RNAP and each cognate site on the DNA was fixed and set to *E*_B_ = 1, while the parameter *E*_CTD_ was varied to change k^CTD^_off_. We assume the binding energy of the RNAP to the cognate sites on the DNA significantly increases (i.e., more favoring binding) when the number of bound contacts exceeds two. As a result, the escape time of RNAP is not affected by variations in *r_T_* and *E_CTD_*.

The total interaction energy *U*_2_ of the RNAP consists of two contributions: 1) the interaction between pairs of the points representing the non-CTD part of the RNAP, and 2) the energy of interaction between the CTD and the non-CTD part. We therefore have

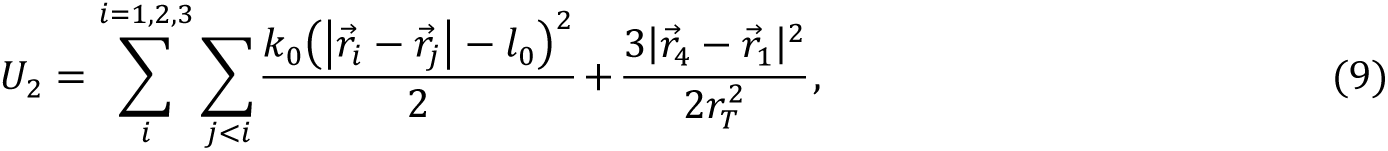

where *k*_0_ = 10 and *r_T_* is the linker length of α-CTD.

When RNAP is close to the DNA, a self-adaptive time step (Δ*t*) is adopted to ensure that 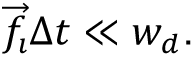

To check that the key conclusions of the Brownian dynamics were not changed by different choices of the parameters described above, we performed a sensitivity analysis where we varied the parameters; the results are described in the SI Appendix.

### Searching for DNA-binding proteins with small, tethered DNA binding domains

To identify proteins that might use the same mechanism for accelerated DNA binding as the one described here for bacterial RNAPs, we searched databases using a two-step filtering process. First, DisProt (https://www.disprot.org/) and UniProt (https://www.uniprot.org/) were both searched for proteins annotated as having both an intrinsically disordered region (IDR) and DNA binding activity. In DisProt, we filtered for proteins with the Intrinsically Disordered Protein Ontology (IDPO) tag “flexible linker/spacer” and the Gene Ontology (GO) tag “DNA binding”. In UniProt, we filtered for proteins that appeared in intrinsically disordered protein databases (DisProt, Intrinsically Disordered proteins with Extensive Annotations and Literature (IDEAL), or Eukaryotic Linear Motif (ELM) resource) and were also annotated for DNA binding. The filtered DisProt and UniProt results were then combined. Second, the combined results were individually screened using information on the proteins’ IDRs from MobiDB (https://mobidb.org/) and on the functions of their structured regions from InterPro (https://www.ebi.ac.uk/interpro). Specifically, we applied the criteria that the protein has two DNA-binding regions linked by a flexible IDR, with the smaller comprising <10% of the residues in the entire protein or >1.5-fold fewer residues than the larger binding region. Overall, a total of 12 proteins (Table S3) met these criteria.

## Supporting information

Full resolution Fig 2C WT off-DNA

Full resolution Fig 2C WT on-DNA

Full resolution Fig 2C zero CTD off-DNA

Full resolution Fig 2C zero CTD on-DNA

Full resolution Fig S2A one CTD off-DNA

Full resolution Fig S2A one CTD on-DNA

Full resolution Fig S2A WT star star off-DNA

Full resolution Fig S2A WT star star on-DNA

## Acknowledgements

A.A. thanks the Clore Center for Biological Physics, Weizmann Institute of Science for its support. This work was supported by grants from NIH (R01 GM081648), Simons Foundation (MPS-SIP-00687713), and The Chan Zuckerberg Initiative.

---

**Figure S1:**
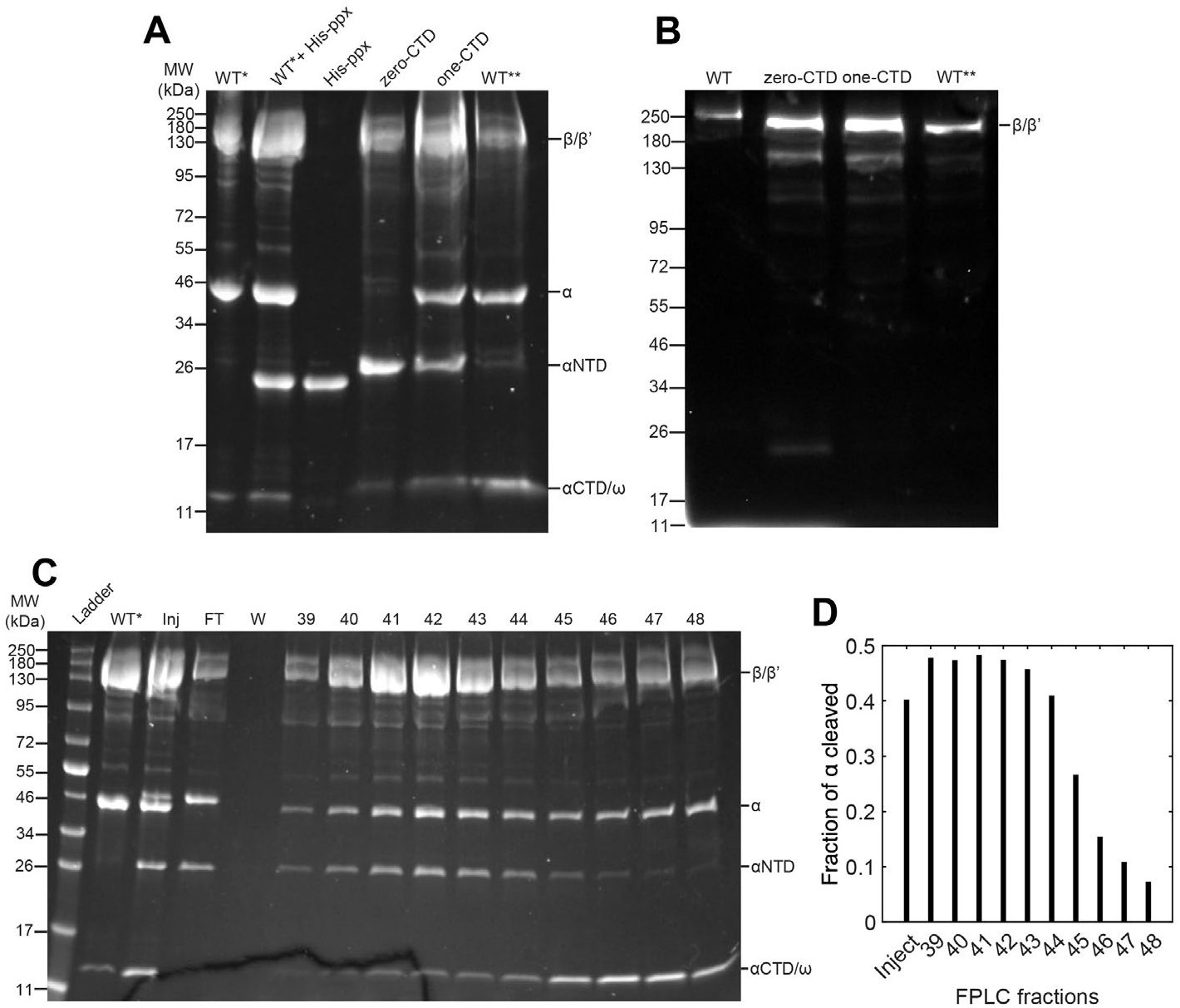
SDS-PAGE electrophoresis gels showing preparation and dye-labeling of RNAP constructs with zero, one, and two CTDs. **(A)** Coomassie Blue stained gel showing WT* RNAP, WT* incompletely digested with his-ppx, his-ppx protease alone, and the zero-CTD, one-CTD, and WT** protein preparations. **(B)** Fluorescence-scanned gel of the indicated protein preparations using DyLight 550 excitation on the ChemiDoc imaging system (Bio-Rad). **(C)** Purification on a His-Trap column (see Methods) of one-CTD RNAP and WT** from a partial cleavage reaction of WT* RNAP with ppx. **(D)** Mole ratio of cleaved to total α subunits for the column fractions in (C), determined by densitometry of the α and α-NTD bands. One-CTD RNAP and WT** RNAP were prepared from fractions 41 and 48, respectively.

**Figure S2:**
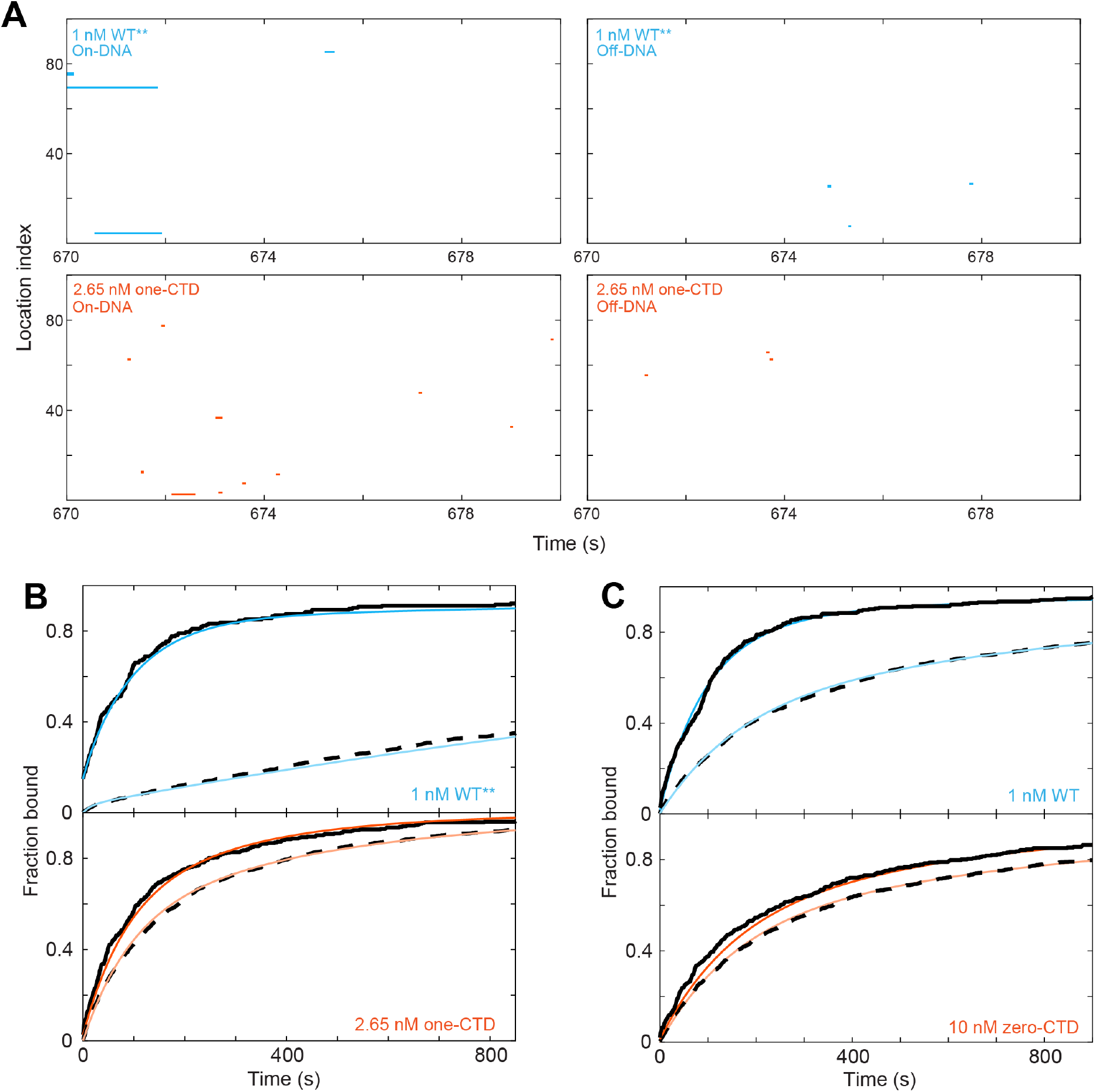
Single-molecule colocalization measurements of the association kinetics of one-CTD RNAP with DNA and an additional replicate of a zero-CTD experiment. **(A)** Rastergram plot of 100 randomly selected DNA or control sites. Each plot shows a magnified view of a 10 s time window in order to display detail; high resolution views of the complete time records are provided as supplemental data files. **(B)** Time-to-first binding measurements for the WT** and one-CTD constructs, plotted and analyzed as in Fig. 2D. **(C)** Time-to-first binding measurements for the WT and zero-CTD constructs (a replicate experiment to that shown in Fig. 2D). Number of observations and fit parameters for (B) and (C) are reported in Table S1.

**Figure S3:**
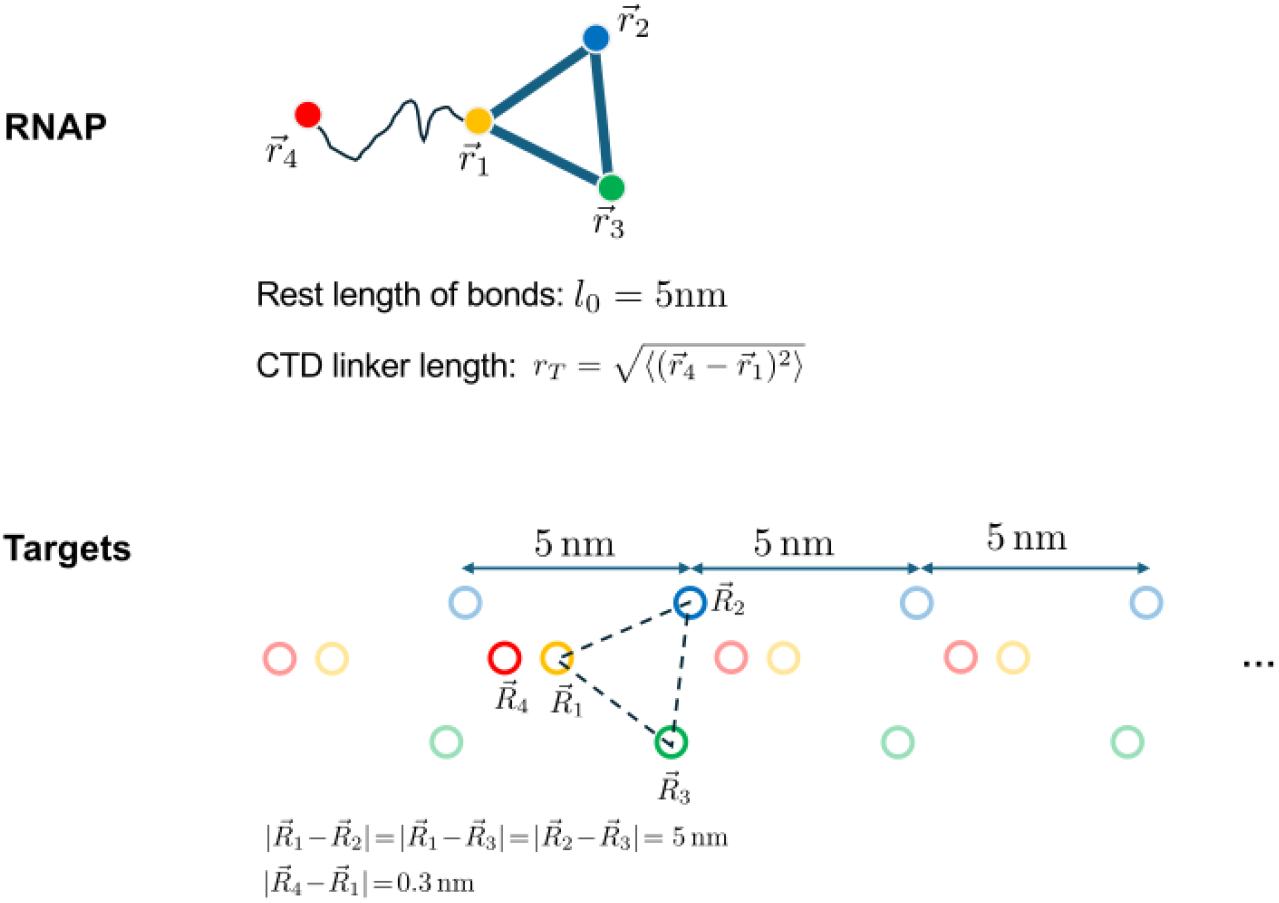
Geometry of RNAP (top) and DNA targets (bottom) used in the Brownian dynamics simulations. The non-CTD part of the RNAP is represented by a triangle with flexible bonds (blue). The CTD (red circle) is a point connected to the triangle at one of the vertices (yellow) via a flexible linker (black line), whose mean end-to-end distance varies in the simulations. All the colored points that make up the RNAP undergo diffusion. The RNAP binds to the targets on the DNA, which are arranged as a periodic repeat of binding sites. For binding to occur the colors of the binding sites on the DNA (red, yellow, blue, green) need to match up to the same-colored sites on the RNAP. Parameters used in the Brownian simulation are summarized in Table S2.

**Figure S4:**
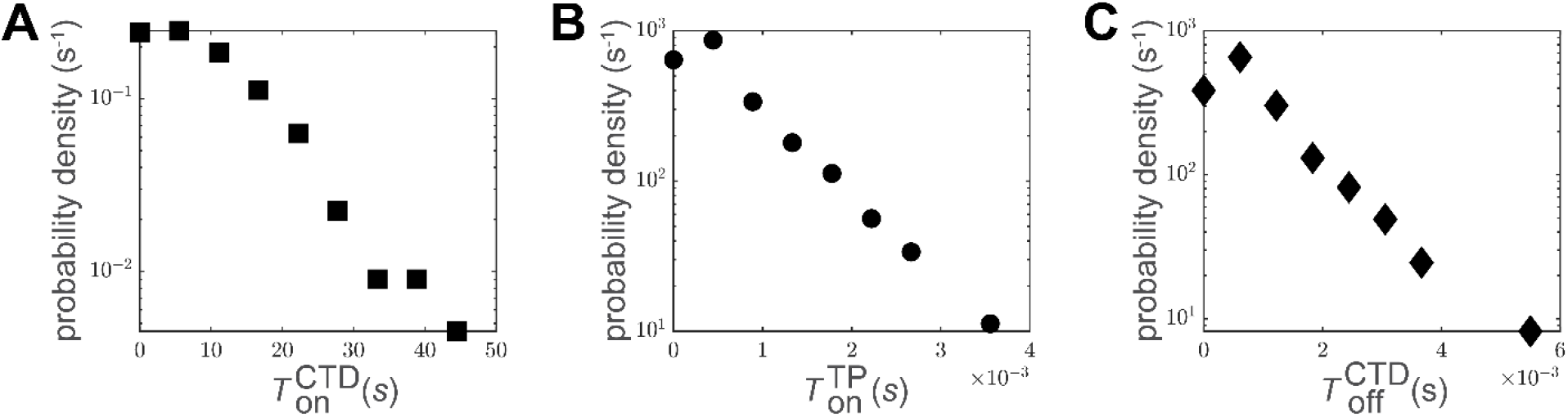
Distribution of times to go between different states, obtained from Brownian dynamics. **(A)** Distribution of times (*T*^CTD^_on_) to go from the unbound state of the RNAP to the intermediate state, in which CTD is bound to the DNA. **(B)** Distribution of times (*T*^TP^_on_) to go from the intermediate state to the bound state, in which the non-CTD part of the polymerase is bound to the DNA. **(C)** Distribution of times (*T*^CTD^_off_) to go from the intermediate state to the unbound state. Parameters used in the Brownian dynamics simulations were *E_B_* = 1 *k_B_T*, *E_CTD_* = 15 *k_B_T* and *r_T_* = 1 nm.

**Table S1:** Fit parameters for measurements of RNAP association kinetics with DNA.

| Figure | RNAP construct | $A_f$ | $k_{on}$<br>($\times 10^{-3} \text{ s}^{-1}$) | $k'_{on}$<br>($\times 10^6 \text{ M}^{-1} \text{ s}^{-1}$) | $a_{ns}$ | $k_{ns,1}$<br>($\times 10^{-3} \text{ s}^{-1}$) | $k_{ns,2}$<br>($\times 10^{-3} \text{ s}^{-1}$) |
| --- | --- | --- | --- | --- | --- | --- | --- |
| Fig. 2D | 1 nM WT | $0.80 \pm 0.05$ | $6.7 \pm 1.1$<br>($N = 381$) | $6.7 \pm 1.1$ | $0.31 \pm 0.08$ | $4.5 \pm 1.1$<br>($N_n = 2,724$) | $1.1 \pm 0.1$ |
| | 25 nM zero-CTD | | $8.8 \pm 2.4$<br>($N = 321$) | $0.35 \pm 0.09$ | $0.59 \pm 0.09$ | $13 \pm 2.6$<br>($N_n = 1,499$) | $4.0 \pm 0.5$ |
| Fig. S2C | 1 nM WT | $0.77 \pm 0.06$ | $7.6 \pm 1.2$<br>($N = 295$) | $7.6 \pm 1.2$ | $0.50 \pm 0.04$ | $4.9 \pm 0.6$<br>($N_n = 1,909$) | $0.8 \pm 0.1$ |
| | 10 nM zero-CTD | | $0.6 \pm 0.5$<br>($N = 361$) | $0.06 \pm 0.05$ | $0.56 \pm 0.05$ | $5.0 \pm 0.6$<br>($N_n = 948$) | $0.9 \pm 0.1$ |
| Fig. S2B | 1 nM WT** | $0.84 \pm 0.04$ | $9.5 \pm 1.2$<br>($N = 215$) | $9.5 \pm 1.2$ | $0.03 \pm 0.17$ | $48 \pm 32$<br>($N_n = 1,428$) | $0.4 \pm 0.1$ |
| | 2.65 nM one-CTD | | $2.4 \pm 1.2$<br>($N = 353$) | $0.91 \pm 0.45$ | $0.57 \pm 0.06$ | $7.6 \pm 1.1$<br>($N_n = 1,479$) | $2.0 \pm 0.2$ |

\* $k_{on}$ and $k_{ns}$ are the apparent first-order rate constants for RNAP binding to DNA and for RNAP non-specific binding to the chamber surface. $k'_{on}$ is the second-order rate constant for the binding to DNA. $A_f$ is the active fraction of DNA molecules capable of RNAP binding. $a_{ns}$ is the relative amplitude between the fast and slow components of the bi-exponential fit to the non-specific binding. The time resolution for all experiments was 65 ms.

**Table S2:** Values of parameters used in the Brownian simulation. Parameter values used in equations 7-9 of the main text. The diffusion coefficient of the center of mass of the triangle representing the non-CTD part of the RNAP (see Figure S2) is 1/3 of the diffusion coefficient of each of the three points of the triangle.

|  |  |
| --- | --- |
| Target size: $w_d$ | 0.5 nm |
| RNAP length: $l_0$ | 5 nm |
| RNAP targets distance | 5 nm |
| Radius of the sphere for search: $R$ | 500 nm |
| Stiffness of the RNAP bond: $k_0$ | $10 k_B T / \text{nm}^2$ |
| RMSD of α-CTD: $r_T$ | 1,2,3,4,5,6,7,8,10,15,25 nm in Fig. 5C |
| Total number of periods | 20 |
| Periodic spacing of RNAP targets | 5 nm |
| Depth of RNAP target potential: $E_B$ | $1 k_B T$ |
| Depth of the guide potential: $E_{CTD}$ | 8,9,10,11,13,15,17,19,21 $k_B T$ in Fig. 5B |
| RNAP diffusion coefficient | $D_0/3 = 0.3 \mu\text{m}^2/\text{s}$ |

**Table S3:** DNA-binding proteins with small tethered DNA binding domains.

| Protein (species) | UniProt ID | Domain organization |
| --- | --- | --- |
| Bloom Syndrome Protein (Human) | P54132 | Within its helicase domain, has two parts that bind DNA where one is 6% of the protein, the other is 39%, and they are connected by a flexible tether |
| DNA-binding protein H-NS (E. Coli) | P0ACF8 | Contains 1 DNA binding region (20% of protein) flexibly linked to another DNA binding region (40% of protein) |
| Homeobox Protein Hox-B1 (Human) | P14651 | Contains two DNA binding regions (5% and 20% of protein) that are flexibly linked |
| Homeobox Protein Hox-B3 (Human) | P14653 | Contains two DNA binding regions (3% and 20% of protein) that are flexibly linked |
| Nucleolar Transcription Factor 1 (Human) | P17480 | Contains a large DNA binding region (37% of protein) flexibly linked to a smaller DNA binding region (6%) which is also flexibly linked to a small DNA binding region (8%) |
| Nucleolar Transcription Factor 1 (Mouse) | P25976 | Contains a large DNA binding region (35% of protein) flexibly linked to a smaller DNA binding region (6%) which is also flexibly linked to a small DNA binding region (9%) |
| Paired Box Protein Pax-6 (Human) | P26367 | Smaller DNA binding region (20% of protein) flexibly linked to larger region (33%) containing multiple DNA binding regions |
| Paired Box Protein Pax-6 (Zebrafish) | P26630 | Smaller DNA binding region (15% of protein) flexibly linked to larger region (32%) containing multiple DNA binding regions |
| Replication Protein A 70 kDa DNA-binding Subunit (Human) | P27694 | Possible DNA binding region (listed as nucleic acid binding protein; 20% of protein) flexibly linked to large region (70%) containing multiple DNA binding regions |
| ETS Domain containing Protein Elk-4 (Human) | P28324 | Small DNA binding region (4% of protein) flexibly linked to larger DNA binding region (21%) |
| Replication Protein A 70 kDa DNA-binding Subunit (Rice) | Q10Q08 | Possible DNA binding region (listed as nucleic acid binding protein; 15% of protein) flexibly linked to large region (70%) containing multiple DNA binding regions |
| Replication Protein A 70 kDa DNA-binding Subunit (Mouse-ear Cress; a plant) | Q9SKI4 | Possible DNA binding region (listed as nucleic acid binding protein; 16% of protein) flexibly linked to large region (69%) containing multiple DNA binding regions |

## Supporting information

Rivera et al., “Bacterial RNA polymerase exemplifies a general physical mechanism for accelerating protein-DNA association”

SI Figures S1 – S4

SI Tables S1 – S3

SI Appendix text and accompanying Figures A1 – A3

## Supplementary data files

Full resolution data for the rastergrams in Fig. 2C and SI Figure S2A

## SI APPENDIX

### Two-tether model of RNA polymerase

In the main text, both in the kinetic model and the Brownian dynamics simulations, we substituted the two alpha-CTDs and their linkers to the two alpha subunits of the RNA polymerase with a single flexible tether with a (weak) DNA binding domain at its end. Here we show that a more detailed kinetic model, which considers the two flexible tethers, leads to the same formula for the speed-up factor derived in the main text. We verify this conclusion using Brownian dynamics that incorporate two tethers.

### Kinetic model

With two flexible tethers the kinetic model depicted in Fig. 4A acquires an additional state (Fig. A1). Now we can distinguish between two intermediate states, one in which a single tether is bound (S_1_) and one in which both tethers are bound (S_2_), both while the non-CTD part of the RNA polymerase is not bound to the DNA. The stably bound state (F) now has two CTDs bound to the DNA in addition to the binding of the non-CTD part of the RNA polymerase.

**Figure A1:**
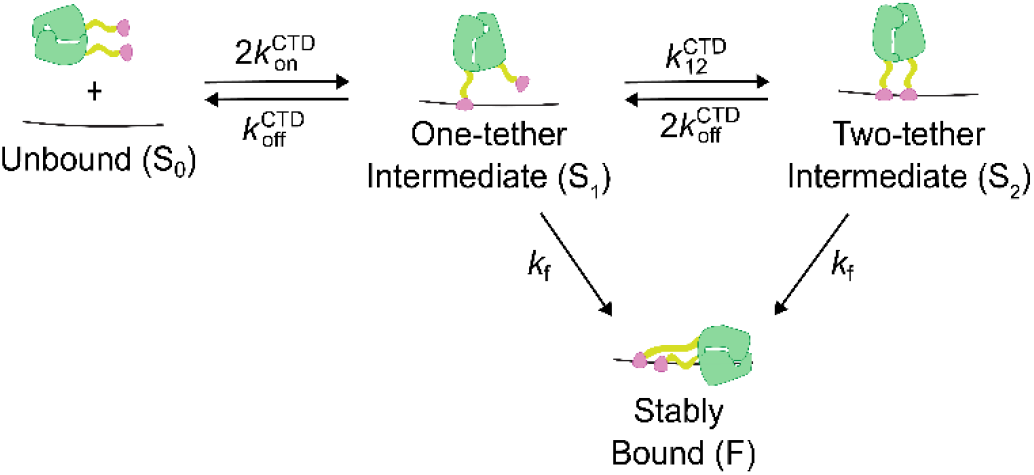
Two-tether model of RNAP binding to DNA. RNAP binding is represented by a Markov process, with four states. Rate constants characterize the transitions between states. S_0_ is the state of RNAP freely diffusing in solution. F is the stably bound state, while S_1_ and S_2_ are intermediate states with one tether or two tethers bound, respectively. To calculate the first passage time from the S_0_ state to the F state (equations A1 and A2) we take the transitions to the F state to be irreversible.

To arrive at a formula that describes the effect of flexible tethers on the rate of RNA polymerase binding to the DNA, we compute the mean first passage time (*T*_0_) from the unbound (U) state to the stably bound, final (F) state. The effective association rate is the reciprocal of this first passage time, *k_eff_* = 1/*T*_0_.

To compute the mean first passage time from state S_0_ to F we make use of the system of linear equations for the mean first passage times, *T*_0_, *T*_1_, and *T*_2_ from the states S_0_, S_1_, and S_2_, respectively, to the final state:

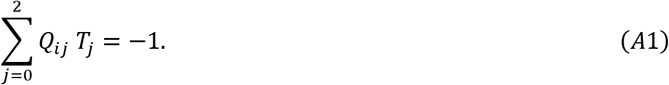

Here *Q* is the transition rate matrix

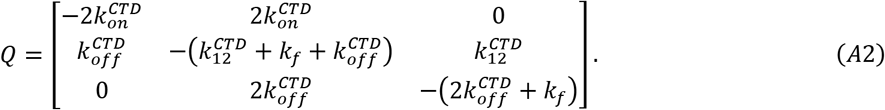

which can be read off the kinetic scheme shown in Fig. A1; the matrix element *Q_ij_* when *i* ≠ *jj* is the rate at which the state S_i_ transitions to state S_j_, while the diagonal elements *Q_ii_* are equal to the negative of the total rate of departure from the S_i_ state.

Solving the system of linear equations for *T*_0_ leads to a rather complicated expression for *k_eff_* = 1/*T*_0_ in terms of all the rate constants:

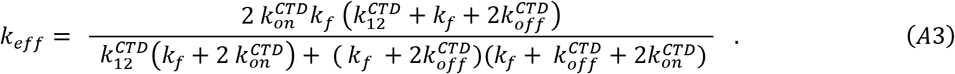

We simplify this equation by assuming that the rate of binding the CTD from solution is much smaller than the rate for it to bind to DNA when the polymerase is tethered by the other CTD, i.e., 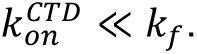. This approximation, which is equivalent to the one made in the main text that led to equation 2, reduces equation A3 to

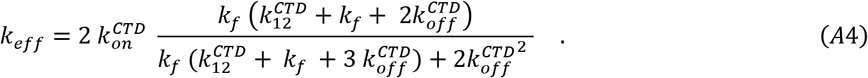

Furthermore, assuming that 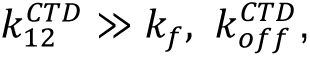, we arrive at an equation which has the exact same form as equation 2 derived for the one-tether case, (dropping the much smaller rate of binding of the polymerase directly from solution, *k^p^_on_*)

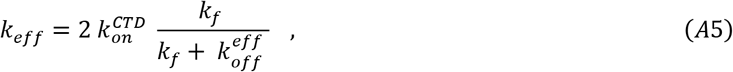

where 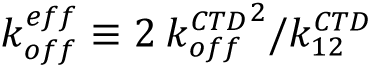 is the effective rate for the CTD to dissociate from the DNA. This functional form makes intuitive sense since the square of the off rate for a single CTD corresponds to double the binding energy of two CTDs when compared to the binding energy of one CTD. The equivalence of the functional form used in the main text, further justifies using the simpler one-tether model for the RNA polymerase.

### Brownian dynamics of two-tether model of RNAP

In the main text we replaced the two alpha-CTDs and their flexible linker-domains with a single polymer. Here we consider a more detailed model of the alpha-CTDs which are described by two polymers connected to the non-CTD part of the polymerase. The non-CTD part is described by a triangle as in the main text, while the two tethers are attached to the centers of two sides of the triangle (Figure A2, inset). All the other parameters that define the Brownian motion of the RNAP and its interactions with the DNA are modeled as in the case of one tether (see Methods). The results for the speed-up of the binding as a function of the off rate for the CTD-DNA complex are shown in Figure A2 for the one tether and two tether cases. While the speed-up is greater in the two-tether case, it follows the same trend as the one tether case.

**Figure A2:**
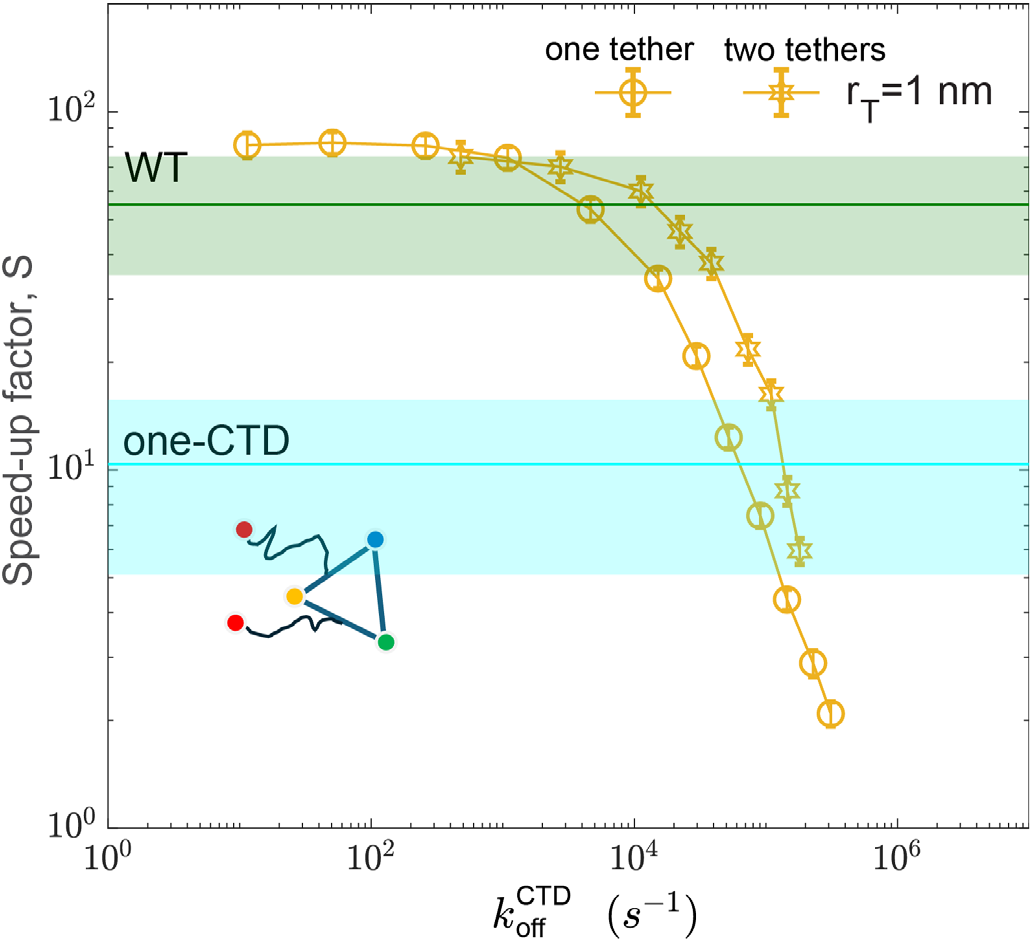
**The speed-up factor as a function of the dissociation rate *k^CTD^_off_*** Comparison of the speed-up factor from Brownian dynamics simulations of the one-tether and two-tether model of RNAP. The one-tether model is same as in the main text, and the two-tether model of RNAP is shown in the inset. In both cases the length of the tether(s) was set to *r_T_* = 1 nm.

### Sensitivity Analysis for Brownian Dynamics Simulations

Here we test the sensitivity of the Brownian dynamics results on the choice of RNAP model. We varied the placement of the flexible tether in the model of the RNAP, as shown in Figure A3A-C. Also, we varied the length of the flexible tether (Figure A3D), and the off rate for the CTD-DNA complex (Figure A3E). While the quantitative results change, the main conclusions we derive from the model, specifically that the transient binding of the alpha-CTD can increase the binding rate by almost two orders of magnitude, remains unchanged.

**Figure A3:**
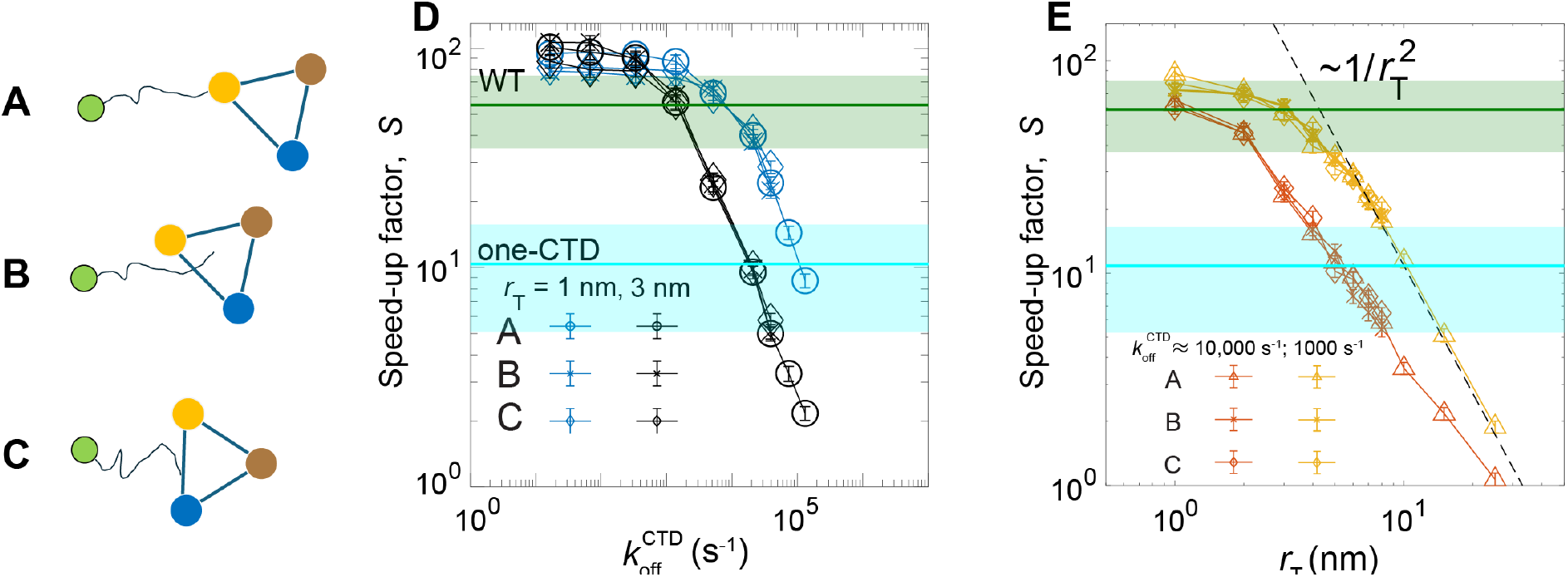
Brownian dynamics results are independent of the tether attachment point on the RNAP. (A, B, **C)** Schematics of the different attachment points of the tether to the non-CTD portion of the polymerase. The CTD (green circle) is tethered to one of the vertices (yellow circle) of the triangle (A), representing the non-CTD part of the polymerase. The tether attachment point can also be at the center of the triangle (B), or at the center of one of the sides of the triangle(C). **(D)** The speed-up factor for two different tether lengths (*r*_T_) as a function of the dissociation rate for the CTD from the DNA (k^CTD^_off_) is independent of the choice of attachment point. **(E)** The speed-up factor for two different CTD dissociation rates as a function of the tether length is independent of the choice of attachment point.

