## Supplementary figures and images for "Bacterial RNA polymerase exemplifies a general physical mechanism for accelerating protein-DNA association"

### Full resolution Fig 2C WT off-DNA

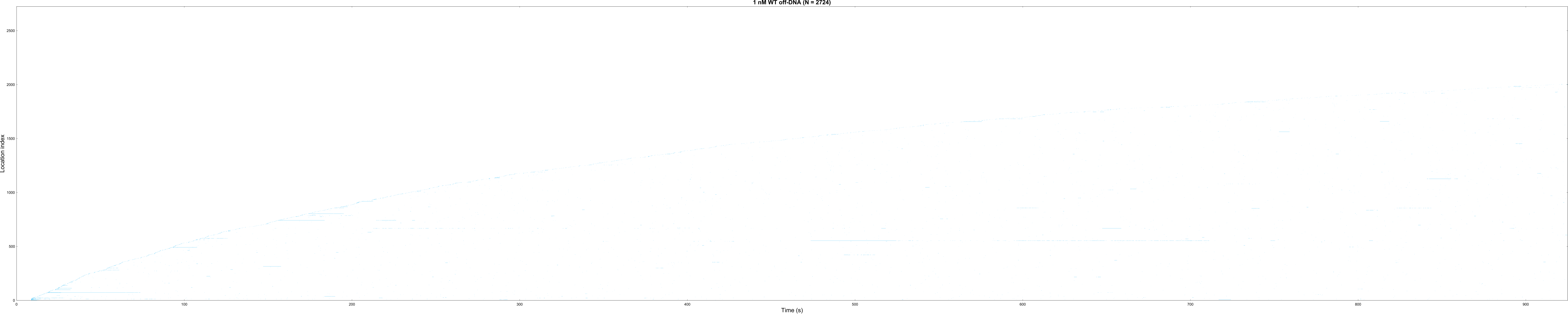

### Full resolution Fig 2C WT on-DNA

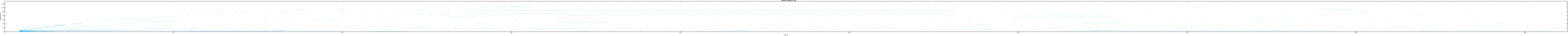

### Full resolution Fig 2C zero CTD off-DNA

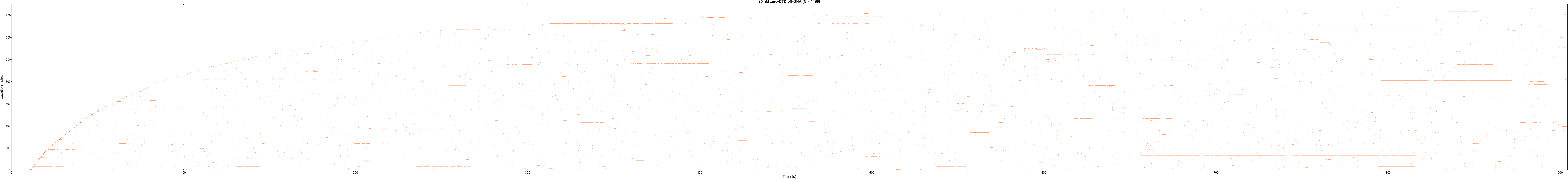

### Full resolution Fig 2C zero CTD on-DNA

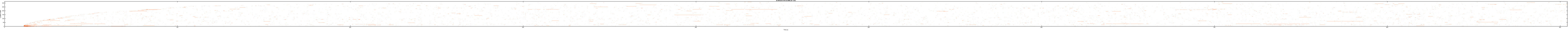

### Full resolution Fig S2A one CTD off-DNA

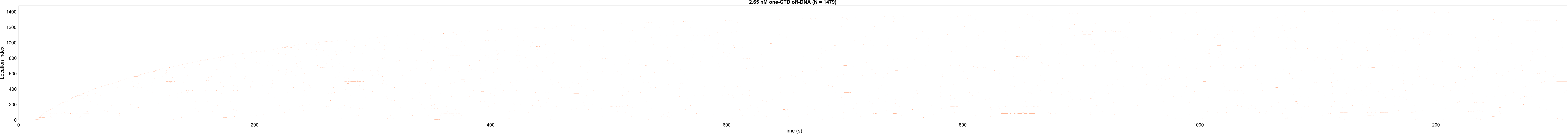

### Full resolution Fig S2A one CTD on-DNA

2.65 nM one-CTD on-DNA (N = 353)

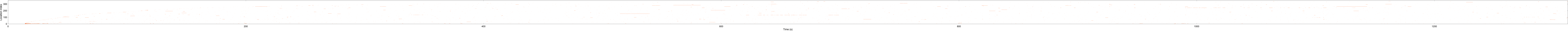

### Full resolution Fig S2A WT star star off-DNA

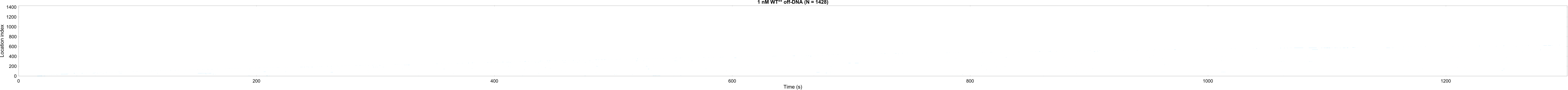

### Full resolution Fig S2A WT star star on-DNA

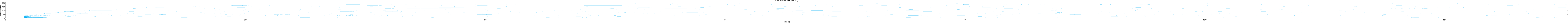
